# All-atom molecular dynamics simulations of synaptic vesicle fusion I: a glimpse at the primed state

**DOI:** 10.1101/2021.12.29.474428

**Authors:** Josep Rizo, Levent Sari, Yifei Qi, Wonpil Im, Milo M. Lin

## Abstract

Synaptic vesicles are primed into a state that is ready for fast neurotransmitter release upon Ca^2+^-binding to synaptotagmin-1. This state likely includes trans-SNARE complexes between the vesicle and plasma membranes that are bound to synaptotagmin-1 and complexins. However, the nature of this state and the steps leading to membrane fusion are unclear, in part because of the difficulty of studying this dynamic process experimentally. To shed light into these questions, we performed all-atom molecular dynamics simulations of systems containing trans-SNARE complexes between two flat bilayers or a vesicle and a flat bilayer with or without fragments of synaptotagmin-1 and/or complexin-1. Our results help visualize potential states of the release machinery en route to fusion, and suggest mechanistic features that may control the speed of release. In particular, the simulations suggest that the primed state contains almost fully assembled trans-SNARE complexes bound to the synaptotagmin-1 C_2_B domain and complexin-1 in a spring-loaded configuration where interactions of the C_2_B domain with the plasma membrane orient complexin-1 toward the vesicle, avoiding premature membrane merger but keeping the system ready for fast fusion upon Ca^2+^ influx.

## Introduction

The release of neurotransmitters by Ca^2+^-triggered synaptic vesicle exocytosis is key for communication between neurons. The high speed of this process arises in part because synaptic vesicles are first tethered at the plasma membrane and undergo a priming process that leaves them ready for fast fusion when an action potential induces Ca^2+^ influx (Sudhof, 2013). Research for over three decades has led to extensive characterization of the machinery that controls neurotransmitter release (Brunger et al., 2018; Rizo, 2018), allowing reconstitution of basic steps leading to synaptic vesicle fusion with the central components of this machinery (Lai et al., 2017; Liu et al., 2016; Ma et al., 2013) and uncovering key aspects of the underlying mechanisms. The SNARE proteins syntaxin-1, SNAP-25 and synaptobrevin form a tight four-helix bundle called the SNARE complex that assembles (zippers) from the N- to the C-terminus to bring the vesicle and plasma membranes together, and is key for membrane fusion (Hanson et al., 1997b; Poirier et al., 1998; Sollner et al., 1993; Sutton et al., 1998). This complex is disassembled by NSF and SNAPs (no relation to SNAP-25) to recycle the SNAREs for another round of fusion (Mayer et al., 1996; Sollner et al., 1993). Munc18-1 and Munc13-1 play central roles in synaptic vesicle priming by organizing assembly of the SNARE complex via an NSF-SNAP-resistant pathway (Ma et al., 2013; Prinslow et al., 2019) whereby Munc18-1 first binds to a self-inhibited ‘closed’ conformation of syntaxin-1 (Dulubova et al., 1999; Misura et al., 2000) and later forms a template for SNARE complex assembly (Baker et al., 2015; Jiao et al., 2018; Parisotto et al., 2014; Sitarska et al., 2017), while Munc13-1 bridges the vesicle and plasma membrane (Quade et al., 2019; Xu et al., 2017) and opens syntaxin-1 (Ma et al., 2011; Yang et al., 2015). Synaptotagmin-1 acts as the Ca^2+^ sensor that triggers fast release (Fernandez-Chacon et al., 2001) in a tight interplay with complexins whereby both synaptotagmin-1 and complexins play inhibitory and active roles (Giraudo et al., 2006; Reim et al., 2001; Schaub et al., 2006; Tang et al., 2006).

Despite this wealth of knowledge, fundamental questions remain about the last steps that lead to fast Ca^2+^-triggered membrane fusion. While there is no doubt that the SNAREs play a central role in this process, it is still unclear how SNARE complex formation induces membrane fusion (Rizo, 2018). Synaptotagmin-1 acts as a Ca^2+^ sensor through its two C_2_ domains (C_2_A and C_2_B), which form most of its cytoplasmic region and bind multiple Ca^2+^ ions via loops at the tip of β-sandwich structures (Fernandez et al., 2001; Sutton et al., 1995; Ubach et al., 1998). These loops are also involved in Ca^2+^-dependent binding of both C_2_ domains to phospholipids, which is critical for release (Fernandez-Chacon et al., 2001; Rhee et al., 2005). The C_2_B domain also binds to PIP_2_ through a polybasic region on the side of the β-sandwich (Bai et al., 2004), which is believed to induce binding to the plasma membrane. Moreover, the C_2_B domain can bind to the SNARE complex through three different surfaces (Brewer et al., 2015; Zhou et al., 2015; Zhou et al., 2017), although only binding through a so-called primary interface (Zhou et al., 2015) is firmly established as physiologically relevant (Guan et al., 2017; Voleti et al., 2020). This interaction is likely important for SNARE complex assembly (Prinslow et al., 2019), thereby facilitating vesicle priming (Zhou et al., 2017), but may inhibit premature fusion. Indeed, recent data suggested that Ca^2+^ induces a tight, PIP_2_-dependent interaction of the C_2_B domain with the membrane, dissociating synaptotagmin-1 from the SNARE complex (Voleti et al., 2020). Synaptotagmin-1 may facilitate fusion by bridging the two membranes (Arac et al., 2006; van den et al., 2011) and/or by inducing membrane curvature (Arac et al., 2006; Martens et al., 2007), but the precise mechanism remains unknown.

Complexin-1 binds tightly to the SNARE complex through a central α-helix that is preceded by an accessory helix (Chen et al., 2002) and may play a stimulatory role in release by promoting formation of a primed state with enhanced release probability (Chen et al., 2002), by protecting trans-SNARE complexes from disassembly by NSF and αSNAP (Prinslow et al., 2019) and/or by synchronizing Ca^2+^-triggered fusion mediated by the SNAREs and synaptotagmin-1 (Diao et al., 2012). The complexin-1 accessory helix inhibits release (Xue et al., 2007), likely because it causes steric clashes with the vesicle, hindering C-terminal assembly of the SNARE complex (Radoff et al., 2014; Trimbuch et al., 2014). These and other findings suggested that complexin-1 and synaptotagmin-1 bind simultaneously to the SNARE complex in the primed state of synaptic vesicles, stabilizing this state and preventing premature fusion; in this model, Ca^2+^ influx relieves the inhibition by inducing dissociation of synaptotagmin-1 from the SNAREs and enables cooperation between synaptotagmin-1 and the SNAREs in promoting fusion (Voleti et al., 2020). However, these various models have not yet been validated.

Major hurdles to elucidate the last steps that lead to fast, Ca^2+^-triggered membrane fusion are the dynamic nature of this process and the fact that the protein complexes that trigger fusion are assembled between two membranes. Although important clues on the nature of the primed macromolecular assembly have been obtained with structural studies of soluble proteins or complexes anchored on one membrane (Chen et al., 2002; Grushin et al., 2019; Voleti et al., 2020; Zhou et al., 2015), this assembly is most likely affected by its location between two membranes. This feature strongly hinders the possibility of crystallization, while application of NMR spectroscopy to determine the structure of the primed assembly is hampered by the large size of any reconstituted two-membrane system and problems with sample stability (Voleti et al., 2021). Conversely, the small size of the SNAREs, synaptotagmin-1 and complexin-1 hinders visualization by cryo-EM. Moreover, it is extremely challenging to capture transient states formed during the pathway to Ca^2+^-triggered membrane fusion experimentally.

Molecular dynamics (MD) simulations offer a powerful tool to analyze dynamic biomolecular processes, and are increasingly being applied to realistic models of cellular membranes, including coarse-graining approaches to extend the simulation time (Marrink et al., 2019). Coarse-grained MD simulations of SNARE-mediated membrane fusion have already been performed (Risselada et al., 2011; Sharma and Lindau, 2018), although the significance of these simulations is uncertain because the continuous yet highly strained helices assumed to be formed by synaptobrevin and syntaxin-1 throughout these simulations are unrealistic from an energetic point of view (see below). Moreover, all-atom simulations are better suited to reproduce the finely-balanced network of interactions between proteins, Ca^2+^ and lipids that are expected to lead to membrane fusion. Although the time scales reachable with all-atom MD simulations are more limited than those attainable with coarse-grained simulations, high performance computing currently allows calculation of microsecond trajectories for systems containing millions of atoms. Indeed, all-atom simulations have already provided important insights into interactions among the components of the primed complex (Bykhovskaia, 2021; Bykhovskaia et al., 2013). Note also that the delay from Ca^2+^ influx into the presynaptic terminal to observation of postsynaptic currents in rat cerebellar synapses at 38°C is 60 μs (Sabatini and Regehr, 1996), and that multiple events occur within this time frame, including Ca^2+^ binding to the sensor, Ca^2+^-evoked synaptic vesicle fusion, opening of the fusion pore, diffusion of neurotransmitters through the synaptic cleft, binding of the neurotransmitters to their postsynaptic receptors and opening of the channels that underlie the postsynaptic currents. These observations suggest that the fusion step occurs in just a few microseconds and therefore that it may be possible to recapitulate Ca^2+^-evoked synaptic vesicle fusion, or at least initial processes that lead to membrane merger, in all-atom MD simulations of this system.

Here we present all-atom MD simulations with explicit water molecules of systems containing four trans-SNARE complexes bridging two flat bilayers or a vesicle and a flat bilayer, without or with fragments of synaptotagmin-1 and/or complexin-1. Because of the limited simulation times, our results cannot lead to definitive conclusions but they help visualize potential trajectories and intermediates along the pathway to fusion and reveal intriguing features, leading to predictions or hypotheses that can be tested experimentally and with additional simulations. Our data indicate that trans-SNARE complexes strongly pull two membranes together, as expected, but have a tendency to induce extended membrane-membrane adhesion interfaces that have been observed experimentally but fuse slowly (Hernandez et al., 2012; Witkowska et al., 2021). Simulations of the primed state of synaptic vesicles lead to a model whereby almost fully assembled trans-SNARE complexes, synaptotagmin-1, complexin-1 and the membranes form a spring-loaded macromolecular assembly that is ready for fast fusion, but steric clashes of complexin-1 with the vesicle hinder the final action of the SNAREs, preventing bilayer adhesion and fusion. Simulations including Ca^2+^ did not lead to membrane fusion but indicate that dissociation of synaptotagmin-1 from the SNARE complex may be a rate limiting step in Ca^2+^-evoked release, and suggest potential mechanisms by which synaptotagmin-1 may cooperate with the SNAREs in inducing membrane fusion.

## Results

### Four trans-SNARE complexes between two flat lipid bilayers

The possibility of observing membrane fusion in the low microsecond time scale in all-atom MD simulations depends critically on the choice of the starting configuration, but the exact nature of the primed state of synaptic vesicles is unknown. Hence, we used the structural and functional information available on this system to generate potential starting configurations. The MD simulations presented here involved systems ranging from 1.7 to 5.9 million atoms. While multiple replicas of each simulation should ideally be carried out to verify the consistency of the results, performing replicated simulations would have limited the number of systems that we could study. Moreover, each simulation included four-trans SNARE complexes bridging two lipid bilayers and the variability in the behavior of the complexes in each simulation already provided insights into the consistency of the observed behaviors. Hence, we chose to use the available high performance computing time to investigate systems with different components and/or distinct geometry, designing each new starting configuration according to what we had learned from the previous simulations. The simulations generated a large amount of data and it is impossible to describe a thorough analysis within the constraints of a single paper. Here we present the main observations from the analyses that we have performed. The files corresponding to the simulations are publically available in Dryad so that interested researchers can perform additional analyses and change the systems as desired to perform additional simulations.

The first system that we built was designed to examine whether SNARE complexes alone can bend two flat lipid bilayers and initiate bilayer fusion. The system contained four trans-SNARE complexes between two square lipid bilayers built with the CHARMM-GUI website (Jo et al., 2008) (https://charmm-gui.org/), which allows a wide choice lipid compositions. The composition of the top bilayer containing anchored synaptobrevin approximated the lipid composition of synaptic vesicles (Takamori et al., 2006) and that of the bottom bilayer with anchored syntaxin-1 was based on the lipid composition of the plasma membrane (Chan et al., 2012) (Table 1). Both bilayers were built with asymmetric lipid distributions in the two leaflets to mimic those present in vivo (Kobayashi and Menon, 2018) (see Methods). The size of both bilayers (26 x 26 nm^2^) was designed to provide space for the SNAREs to bend the membranes and induce fusion while limiting the overall size of the system. The number of SNARE complexes was based on symmetry considerations and the finding that fast vesicle fusion typically observed in synapses requires at least three SNARE complexes (Mohrmann et al., 2010). For simplicity, the SNARE complexes contained the four SNARE motifs, the transmembrane (TM) sequences of syntaxin-1 and synaptobrevin, and the juxtamembrane linkers between their respective SNARE motif and TM region, but did not include the syntaxin-1 N-terminal region or the long linker between the two SNAP-25 SNARE motifs.

**Table 1.**
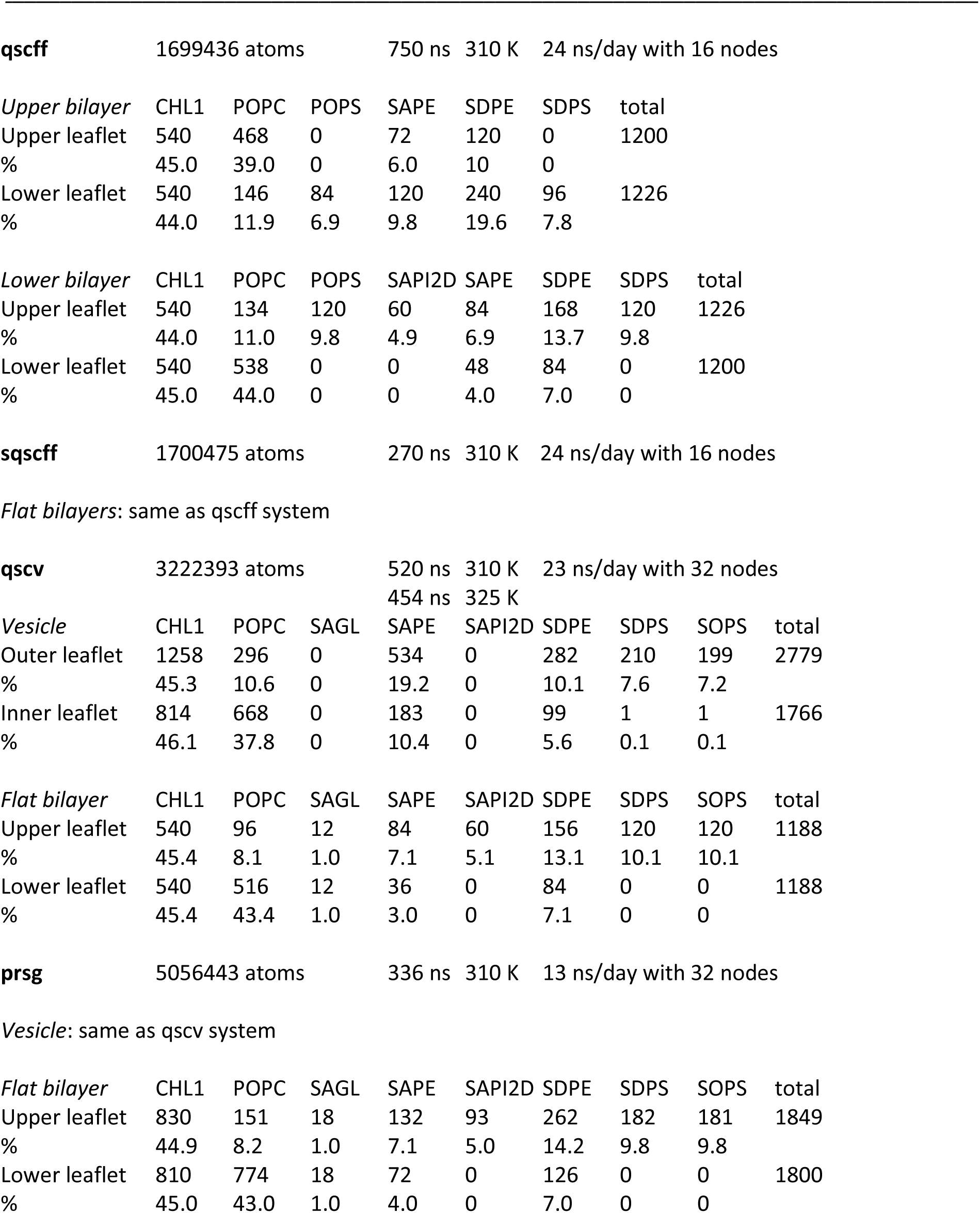
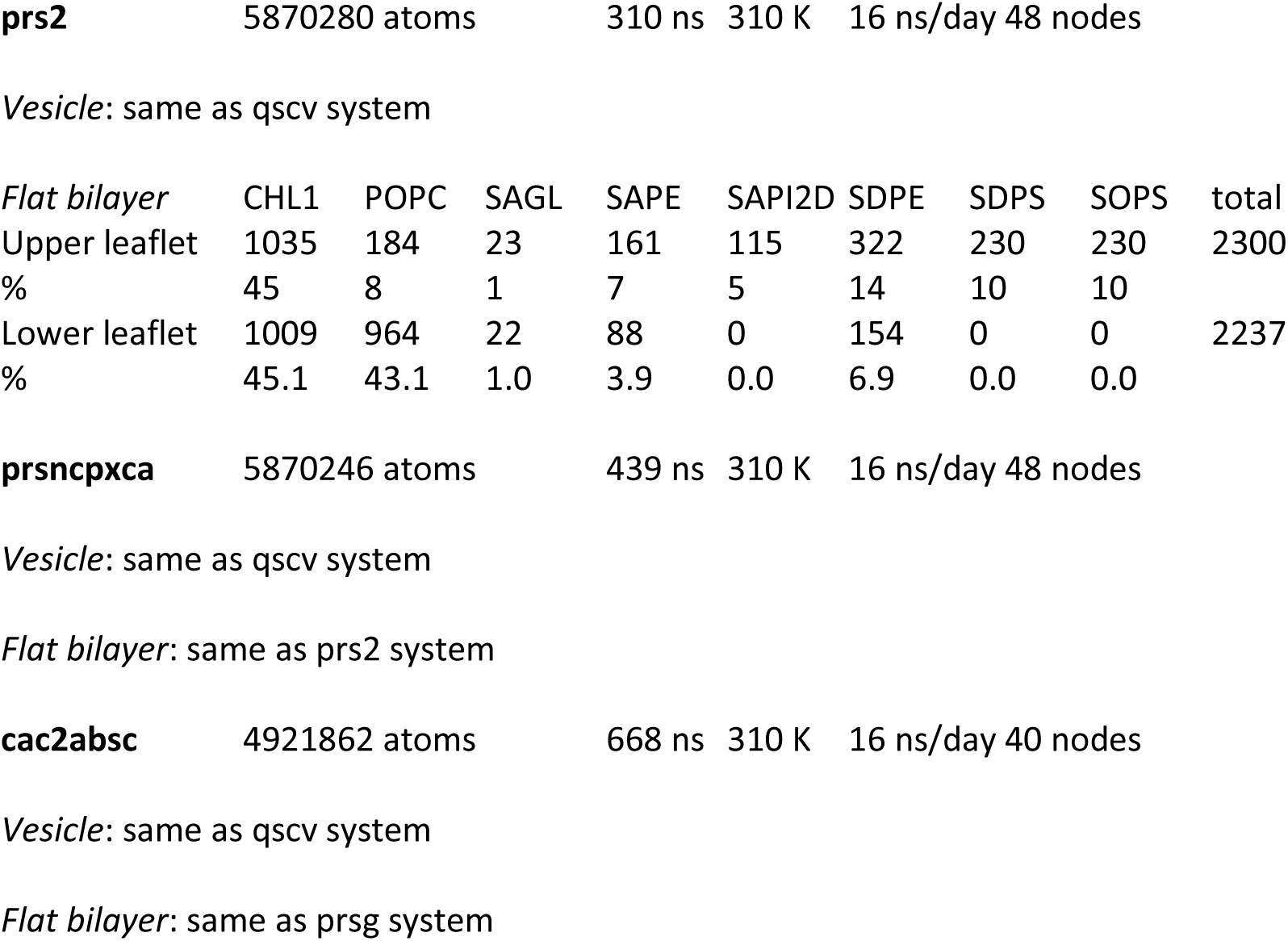
Size in atoms, length of productions MD simulations, temperature, speed of the simulations on Frontera at TACC and lipid composition of the flat bilayers and the vesicle of the different systems

A key aspect in the design of realistic potential states of trans-SNARE complexes is the conformation of the juxtamembrane linkers of syntaxin-1 and synaptobrevin. Popular models of SNARE-mediated membrane fusion depicted continuous helices spanning the SNARE motifs, juxtamembrane linkers and TM regions for both synaptobrevin and syntaxin-1, envisioning that these helices can bend to accommodate the geometry of trans-SNARE complexes (Hanson et al., 1997a; Sutton et al., 1998; Weber et al., 1998). Correspondingly, restraints were used to force such continuous helical conformations in coarse-grained MD simulations, assuming that the helical structures bend with a certain stiffness (Risselada et al., 2011). However, the bending of the helices required to form trans-SNARE complexes leads to unrealistic conformations that are unfavorable energetically, as such bent helices have a distorted geometry and are not commonly observed in protein structures. Thus, the helical restraints might have played a key role in membrane fusion in these simulations. Although continuous helices were observed in the crystal structure of a cis-SNARE complex that represents the configuration occurring after membrane fusion (Stein et al., 2009), the natural expectation is that the helical structure must break somewhere to accommodate the geometry of a trans-SNARE complex, most likely at the juxtamembrane linker. This expectation has been supported experimentally (Kim et al., 2002) and with all-atom MD simulations (Bykhovskaia, 2021). Moreover, helix continuity in the linkers is not required for neurotransmitter release (Kesavan et al., 2007; Zhou et al., 2013). Thus, to generate trans-SNARE complexes for our simulations, we started with the crystal structure of the cis-SNARE complex (adding a few absent residues at the C-termini of syntaxin-1 and SNAP-25) but we did not impose restraints on the conformation of the juxtamembrane linkers. Since the N-terminal half of the SNARE four-helix bundle is more stable than the C-terminal half (Chen et al., 2002; Gao et al., 2012) and is more distal from the membrane, we imposed position restraints for only the N-terminal half. In addition, we used position restraints to force the TM regions of synaptobrevin and syntaxin-1 to designed locations for insertion in their corresponding bilayers.

A short (1 ns) restrained MD simulation in water was sufficient for this purpose and led to unstructured conformations for the juxtamembrane linkers without substantially altering the four-helix bundle even at the C-terminal half, which was not restrained (Figure 1–figure supplement 1A). Four copies of the resulting trans-SNARE complex were generated by translations and rotations (Figure 1–figure supplement 1B), and were merged with the two bilayers to generate the initial configuration of this system (Figure 1A), which contained 1.7 million atoms after solvation. We then carried out an unrestrained production simulation of this system for 750 ns at 310 K. As expected, the two membranes became almost circular to minimize tension and were gradually drawn together by the SNAREs, although the minimal distance between the bilayers reached a plateau (Figure 1–figure supplement 1C,D). The two bilayers were actually drawn to each other on one side first (at about 110 ns, Figure 1B) and later on the other side (Figure 1C), leading to close packing of the lipids against the SNARE four-helix bundles (Figure 1D). It is noteworthy that the four SNARE complexes were zippered at the C-terminus to the same extent as in the initial configuration. Thus, the two membranes were pulled together because extensive interactions were established between the juxtamembrane linkers and the membranes during the simulations. Such interactions were not unexpected, as both linkers contain abundant basic residues, the synaptobrevin linker in addition contains hydrophobic residues, and both linkers were shown to interact with the adjacent membrane (Brewer et al., 2011; Kim et al., 2002). Since much of the SNARE four-helix bundle is negatively charged, these findings suggest that any electrostatic repulsion existing between the SNARE four-helix bundle and the membranes can be readily overcome by the linker-bilayer interactions together with the high stability of the SNARE four-helix bundle. During the 750 ns of the simulation we occasionally observed mild buckling of the syntaxin-1 membrane, but the buckling was reversible and there was no progress toward fusion. These findings suggest that four trans-SNARE complexes are unable to fuse two flat bilayers in the 1 μs time scale.

**Figure 1.**
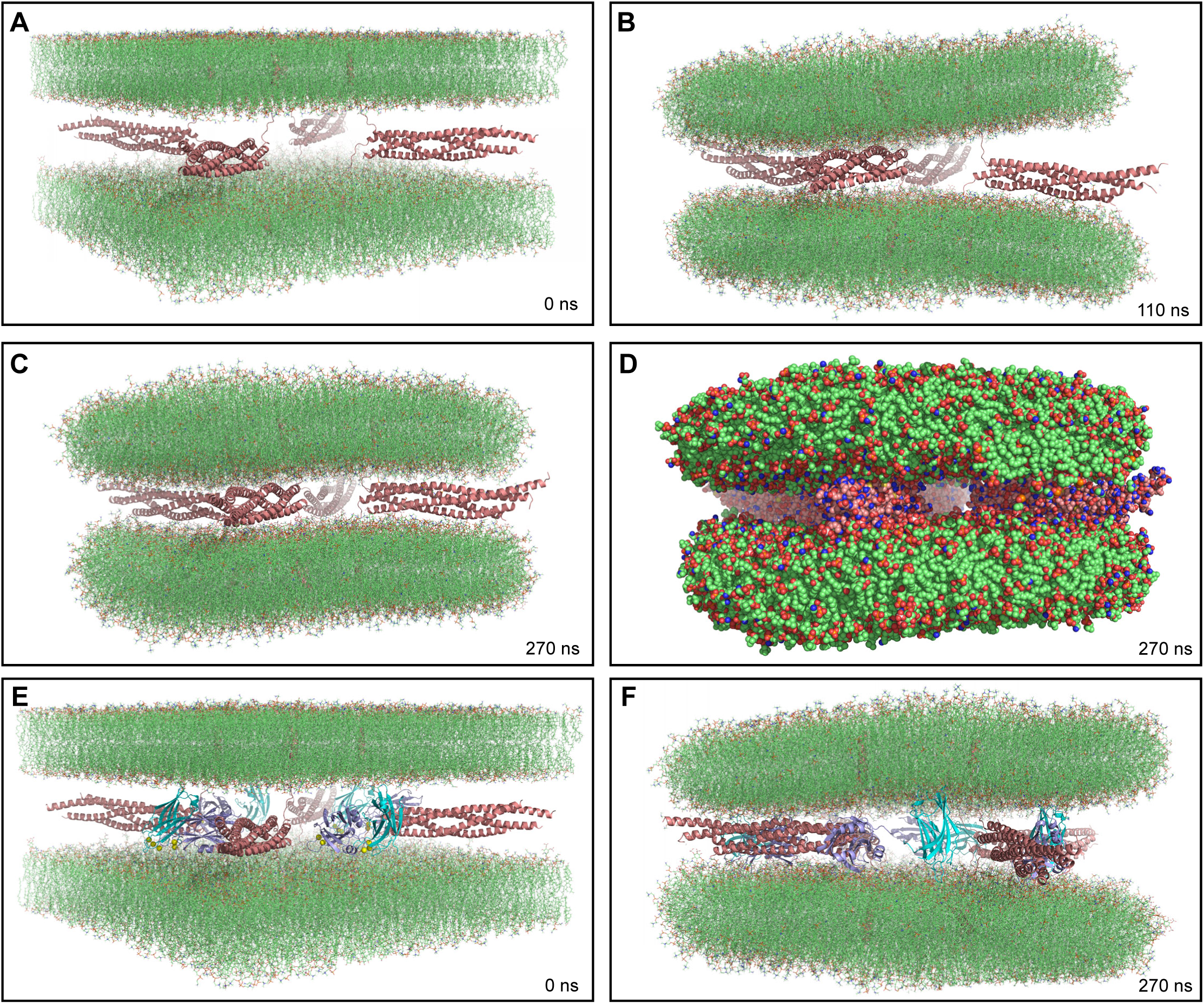
MD simulations of four trans-SNARE complexes bridging two flat bilayers. (**A-C**) Initial configuration of the system with SNARE complexes only (**A**), and snapshots of the MD simulation after 110 and 270 ns (**B, C**). The SNARE complexes are illustrated by ribbon diagrams in salmon. The lipids are shown as thin stick models. (**D**) Snapshot of the same MD simulation at 270 ns showing all non-solvent atoms as spheres. (**E-F**) Initial configuration of the system containing four Ca^2+^-bound synaptotagmin-1 C_2_AB molecules in addition to the four trans-SNARE complexes (**E**) and snapshot of the simulation at 270 ns (**F**). SNARE complexes are illustrated by ribbon diagrams in salmon and the C_2_AB molecules are shown as ribbon diagrams with C_2_A in cyan and C_2_B in violet. The lipids are shown as thin stick models. The atom color code for the lipids is: carbon lime, oxygen red, nitrogen blue, phosphorous orange. Ca^2+^ ions are shown as yellow spheres.

To explore whether synaptotagmin-1 might cooperate with the SNAREs in bending two flat bilayers to initiate membrane fusion, we performed another simulation with an analogous system where we included a fragment spanning the two C_2_ domains of synaptotagmin-1 (C_2_AB) bound to five Ca^2+^ ions (Fernandez et al., 2001; Ubach et al., 1998). A restrained MD simulation of C_2_AB alone was first performed to orient the Ca^2+^-binding loops of both C_2_ domains in similar directions (Figure 1-figure supplement 1E) such that they can readily bind to the same membrane. For copies of the resulting C_2_AB structure were interspersed between the four trans-SNARE complexes but without contacting them (Figure 1E). During a 270 ns production MD simulation of this system, we observed that the C_2_AB molecules hindered the action of the trans-SNARE complexes in bringing the two bilayers closer (Figure 1-figure supplement 1F), particularly when the C_2_ domains bind to one bilayer through the Ca^2+^-binding loops and to the other bilayer via the opposite side of the β-sandwich, which is basic (Figure 1F). Although such bilayer-bilayer bridging might help in fusion (Arac et al., 2006) in a different configuration, it appeared that such potential action would require a much longer time scale in this system and we did not continue this simulation.

### Four trans-SNARE complexes bridging a vesicle and a flat bilayer

Based on the results from the simulations with four trans-SNARE complexes between two flat bilayers, we reasoned that fusion in the low microsecond time scale might require the geometry occurring at synapses, where small synaptic vesicles (ca. 40 nm diameter) fuse with the (approximately flat) plasma membrane. To test this notion, we built a system with four trans-SNARE complexes bridging a vesicle and a flat bilayer (Figure 2A), again mimicking the lipid compositions of synaptic vesicles and the plasma membrane. The four trans-SNARE complexes were slightly modified with respect to those used in the simulations between two flat bilayers, using a restrained MD simulation to tilt the synaptobrevin TM regions such that they were perpendicular to the vesicle surface, and to tilt the SNARE four-helix bundles such that their long axis had similar angles with respect to the vesicle and the flat bilayer (Figure 2B, Figure 2–figure supplement 1A). The flat bilayer was a square of 26 x 26 nm^2^ and matched the diameter of the vesicle, which was 26 nm and was chosen as a compromise between making the system realistic and minimizing the overall size of the system to limit the time required for MD simulations. The vesicle was practically in molecular contact with the flat bilayer so that the system was poised for fusion. Since the lipid density of the vesicle was close to but not optimal, holes appeared in an initial production MD simulation. The holes were filled manually in an iterative process until the vesicle was stable (see Methods). During this procedure, the flat bilayer became circular and the vesicle became slightly smaller (24 nm diameter) (Figure 2–figure supplement 1B,C), but the diameter remained stable in subsequent production runs. The solvated system contained 3.2 million atoms.

**Figure 2.**
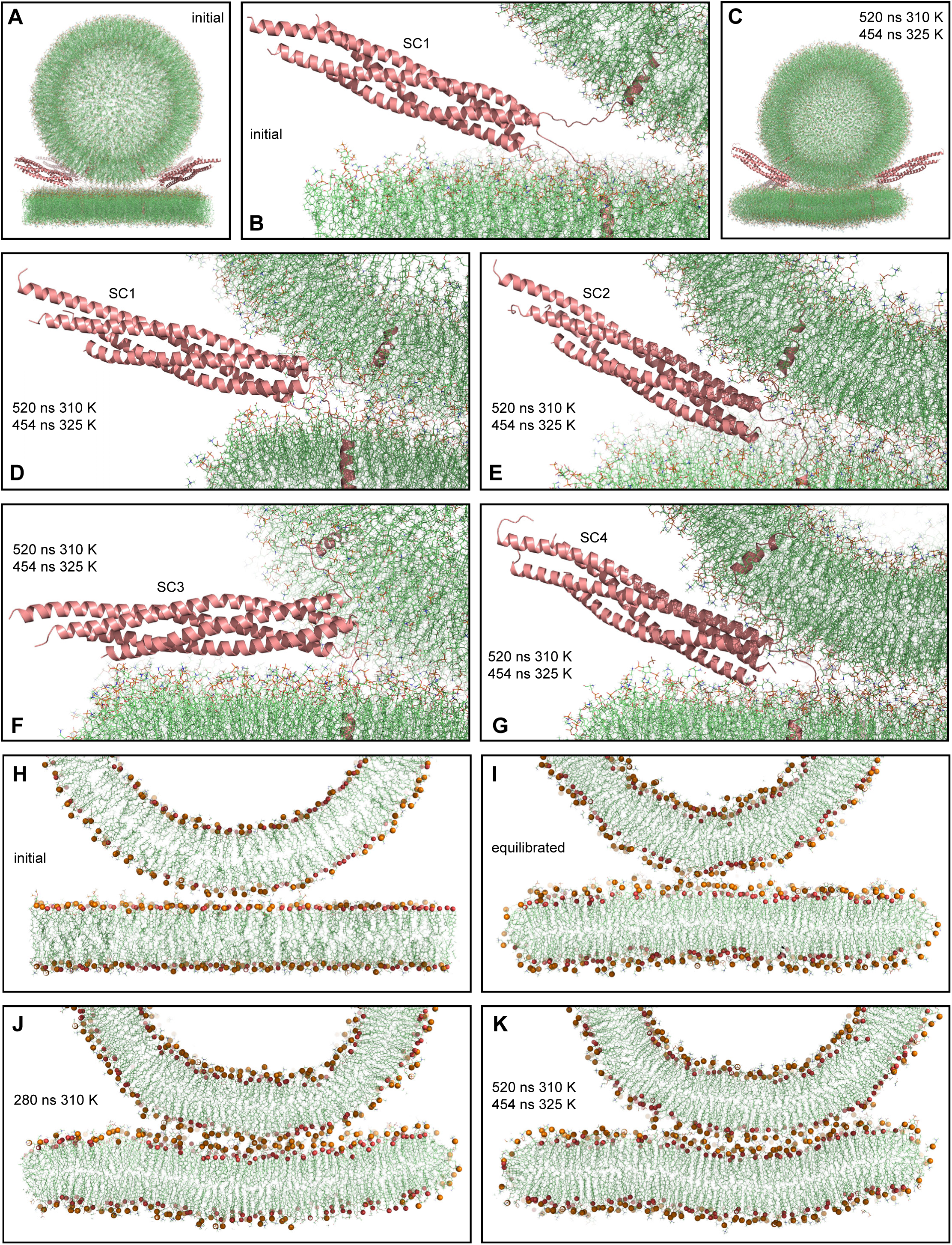
MD simulation of four trans-SNARE complexes bridging a vesicle and a flat bilayer. (**A**) Overall view of the initial system. (**B**) Close up view of one of the trans-SNARE complexes in the initial system. (**C**) Snapshot of the system after a 520 ns MD simulation at 310 K and a 454 ns simulation at 325 K. (**D-G**) Close up views of the four trans-SNARE complexes (named SC1-SC4) after the 520 ns MD simulation at 310 K and the 454 ns simulation at 325 K. In (**A-G**), the SNARE complexes are illustrated by ribbon diagrams in salmon. The lipids are shown as thin stick models (carbon lime, oxygen red, nitrogen blue, phosphorous orange). (**H-K**) Thin slices of the system in its initial configuration (**H**), after the equilibration steps (**I**), after 280 ns at 310K (**J**) and after 520 ns at 310 K and 454 ns at 325 K. In (**H-K**) Phosphorous atoms of phospholipids and the oxygen atoms of cholesterol molecules are shown as spheres to illustrate the approximate locations of lipid head groups.

With the system equilibrated, we performed a production run of 520 ns at 310 K. Although we observed occasional flips of cholesterol molecules, there were no persistent perturbations of the bilayers that might signal the initiation of fusion. We raised the temperature to 325 K and carried out a production run of 454 ns in an attempt to accelerate fusion, but observed similar results. The final configuration illustrates that the vesicle diffused to some extent to one side with respect to the flat bilayer during the simulations (Figure 2C). The four-helix bundles of the four trans-SNARE complexes remained fully assembled up to the last hydrophobic layer [referred to as layer +8 (Sutton et al., 1998)] at the end of the simulations (Figure 2D-G), as in the initial configuration of the system (Figure 2B). One of the SNARE complexes became parallel to the flat bilayer (Figure 2F) whereas the other three had similar orientations as in the starting configuration (Figure 2B,D,E,G). The juxtamembrane linkers of synaptobrevin and syntaxin-1 established extensive interactions with the lipids early in the simulations. These interactions contributed to pull the membranes together and, after 280 ns of the simulation at 310 K, we observed that the bottom of the vesicle was flattened, resulting in an extended interface with the flat bilayer (compare the slice view of Figure 2J with those of the initial configurations in Figure 2H,I). To corroborate these findings quantitatively, we calculated the number of contacts between oxygen atoms of the vesicle and the flat bilayer as a function of time. We assigned a contact to each oxygen-oxygen distance below 1 nm, which is a common cutoff used to calculate van der Waals and electrostatic interactions between atoms. The results showed that the number of contacts increased rapidly up to about 300 ns and then leveled off (Figure 2-figure supplement 1D). The extended interface persisted until the end of the simulation at 325 K, and during this simulation the flat bilayer became slightly curved to adapt to the shape of the vesicle (Figure 2K).

These finding correlates with results obtained in cryo-EM analyses of liposome fusion reactions mediated by the neuronal SNAREs, which revealed extended interfaces between the liposomes that are referred to as tight docking intermediates (Hernandez et al., 2012). These intermediates eventually evolve to yield membrane fusion, but fusion occurs in the second-minute time scale (Hernandez et al., 2012; Witkowska et al., 2021). Hence, it is unlikely that such extended interfaces occur in the pathway that leads to Ca^2+^-triggered synaptic vesicle fusion in microseconds. Note also that the energy required to initiate membrane fusion is expected to increase with the area of the interface between the two membranes, as larger areas require more lipid molecules to be rearranged. Interestingly, cryo-EM images of reconstitution reactions including additional components of the release machinery suggested that these additional components prevent formation of extended interfaces, favoring interfaces with smaller contact area between the bilayers that are referred to as point-of-contact interfaces (Gipson et al., 2017).

### Simulations of the primed state of synaptic vesicles

Overall, our simulations do not rule out the possibility that SNAREs alone might be able to induce membrane fusion in the low microsecond time scale, as it is plausible that other geometries might be more efficient in inducing fusion, for instance if the trans-SNARE complexes were placed closer to each other and to the center of the vesicle-flat bilayer interface. However, the correlation of our results with the cryo-EM images of reconstitution experiments suggests that fast fusion requires other proteins. Formation of a primed state of synaptic vesicles that is ready for fast release is the key to achieve fast Ca^2+^-triggered fusion in synapses. The exact nature of this state is unclear and a recent model that can explain a large amount of data available on presynaptic plasticity actually invokes two primed states: one that involves partially assembled SNARE complexes and has low release probability (referred to as loose state), and another where the SNARE complexes are more fully assembled and has a much higher release probability (referred to as tight state) (Neher and Brose, 2018). Below we use the term primed state to refer to the tight state. Because synaptotagmin-1 and complexins are critical for fast, Ca^2+^-triggered neurotransmitter release (Fernandez-Chacon et al., 2001; Reim et al., 2001), both proteins are most likely bound to the SNARE complex in this primed state. A model of this state (Voleti et al., 2020) was proposed based on crystal structures of the SNARE complex bound to a complexin-1 fragment (Chen et al., 2002) or to the synaptotagmin-1 C_2_B domain through the primary interface (Zhou et al., 2015), as well as a cryo-EM structure of synaptotagmin-1 bound to lipid nanotube-anchored SNARE complex (Grushin et al., 2019). In this model, complexin-1 and the C_2_B domain bind to opposite sides of the SNARE four-helix bundle, and the C_2_B domain binds to the plasma membrane through a polybasic region on the side of the C_2_B domain opposite to the primary interface. However, the orientation of this macromolecular assembly with respect to the vesicle and plasma membranes, and the extent to which the SNARE complex is zippered, are unclear.

To address these questions and gain insights into the nature of the primed state that is ready for fast Ca^2+^-triggered membrane fusion, we built a system with four trans-SNARE complexes bridging a vesicle and a flat bilayer, each bound to a complexin-1 fragment and the synaptotagmin-1 C_2_AB fragment (below referred to as primed complexes). The system was designed to resemble a potential arrangement of the primed state, but implementing some flexibility such that the system could progress towards a preferred configuration of the proteins with respect to the two membranes. To generate the primed system, we used the vesicle described above, after the equilibration procedure, and we built a square flat bilayer of 31.5 x 31.5 nm^2^ to provide sufficient space for interactions with the proteins. The flat bilayer was placed below the vesicle with a minimum separation of 2.3 nm to provide some space for the system to evolve. The relative orientation of the two C_2_ domains in the initial conformation of the C_2_AB fragment was designed to have the C_2_A domain Ca^2+^-binding loops orientated toward the flat bilayer while the C_2_B domain can bind to the flat bilayer through its polybasic face. The complexin-1 fragment included all the residues that were observed in the crystal structure of the complexin-1-SNARE complex (residues 32-72) (Chen et al., 2002) plus five additional N-terminal residues that may be important for the steric clashes of complexin-1 with the vesicle, which were proposed to underlie the inhibitory activity of the accessory helix (Trimbuch et al., 2014). Thus, the complexin-1 fragment spanned residues 27-72 [Cpx1(27-72)].

To create the initial configurations of the primed complexes, we designated four positions to place the SNARE four-helix bundles bound to Cpx1(27-72) and to the synaptotagmin-1 C_2_AB through the primary interface as determined by crystallography (Chen et al., 2002; Zhou et al., 2015). The C_2_B domain polybasic region was placed close to but not contacting the flat bilayer to limit the bias introduced by the initial configurations of the primed complexes. We then performed a 1 ns MD simulation starting with the four trans-SNARE complexes generated for the vesicle-bilayer system (Figure 2–figure supplement 1A) but bound to Cpx1(27-72), and C_2_ABs in their final desired positions, using position restraints to force all heavy atoms to their designed positions, except for atoms from the C-terminal half of the SNARE four-helix bundle and the juxtamembrane regions, which were not restrained. Because the separation between the positions designed for the synaptobrevin and syntaxin-1 TM regions was increased, this procedure led to partial unfolding of the C-terminal halves of the SNARE four-helix bundles but to distinct extents (Figure 3A,B,D,F,H,J, Figure 3-figure supplement 1), thus yielding a variety of starting points. The final system contained 5.1 atoms after solvation. We note that another crystal structure suggested an additional binding mode of the C_2_B domain on the SNARE complex (Zhou et al., 2017), but its significance is currently unclear (Voleti et al., 2020). For simplicity, we did not include additional C_2_AB molecules binding through this mode, but the potential implications of this binding mode in light of our results are described in the discussion section.

**Figure 3.**
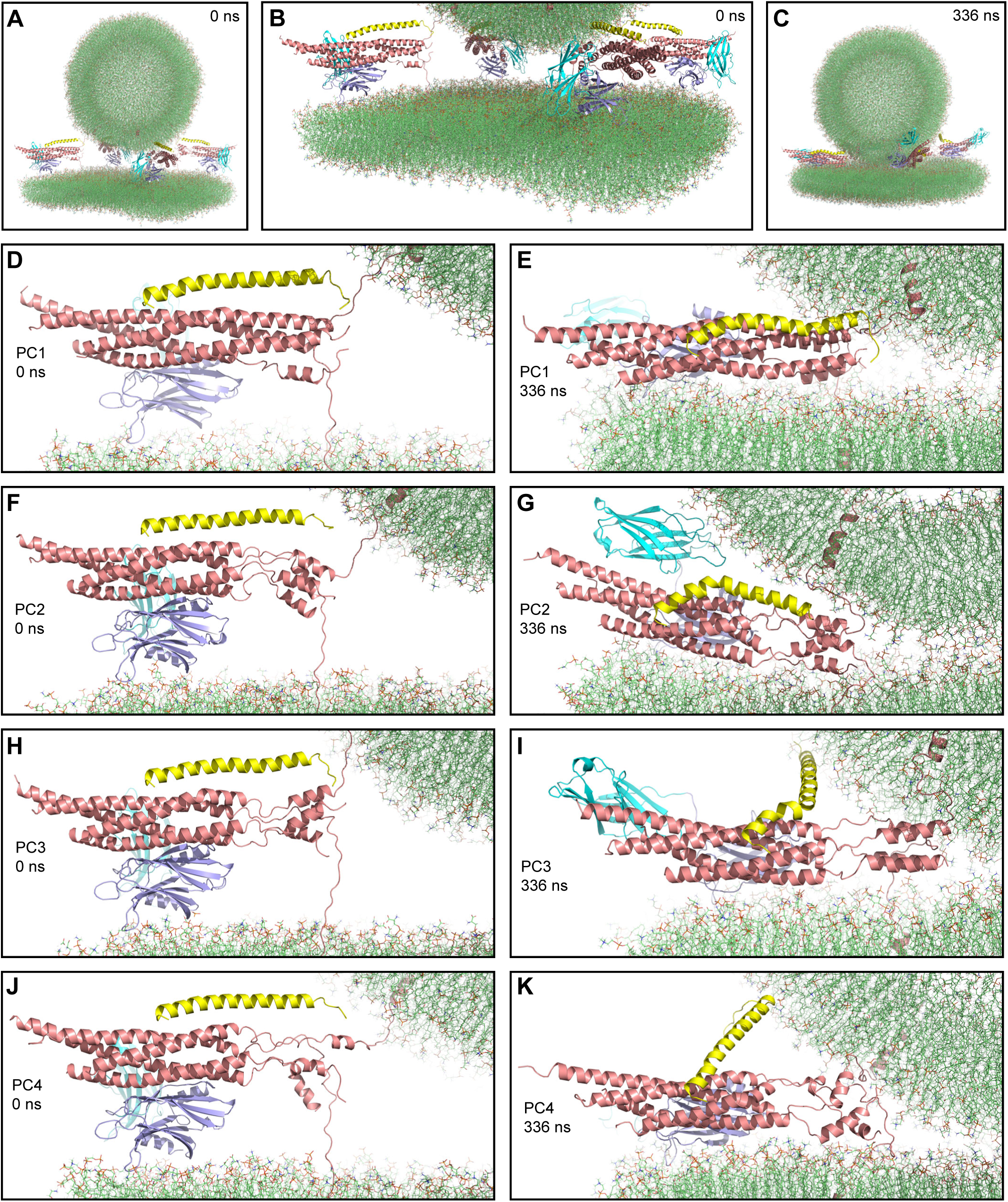
First MD simulation of primed complexes bridging a vesicle and a flat bilayer. (**A**) Overall view of the initial system after equilibration. (**B**) Close up view of the four primed complexes in the initial system after equilibration. (**C**) Snapshot of the system after a 336 ns MD simulation. (**D-K**) Close up views of the individual primed complexes (named PC1-PC4) in the initial configuration (**D,F,H,J**) and after the 336 ns MD simulation (**E,G,I,K**). In all panels, the primed complexes are illustrated by ribbon diagrams, with the SNAREs in salmon, Cpx1(27-72) in yellow and the synaptotagmin-1 C_2_AB fragment in cyan (C_2_A domain) and violet (C_2_B domain). The lipids are shown as thin stick models (carbon lime, oxygen red, nitrogen blue, phosphorous orange).

After equilibration, we carried out a production simulation of 336 ns that resulted in the state shown in Figure 3C. Most substantial changes in the system occurred early in the simulation and each of the primed complexes appeared to reach a stable or metastable configuration by the end. The primed complexes exhibited some common behaviors and also distinct features. The SNARE four-helix bundle of one of the primed complexes (PC1) was almost fully assembled at the start of the simulation (up to layer +7, with a break in a SNAP-25 helix) and remained equally assembled at the end (Figure 3D,E). The C-terminal halves of the SNARE four-helix bundles of the other three complexes were considerably more disrupted and, although they exhibited substantial changes during the simulation, they did not progress toward full assembly (Figure 3F-K). Interestingly, a few of the most C-terminal layers (+5 to +7) still formed a four-helix bundle in two complexes (PC2 and PC3, Figure 3F,G,H,I) and hence they may be particularly stable, but this feature did not seem to facilitate reassembly of the section of the four-helix bundle that was disrupted. Hence, although coil-to-helix transitions are known to occur very fast, in the 100 ns time scale (Munoz and Cerminara, 2016), it appears that the constraints placed on the motions of the SNAREs in this complex system hinder the evolution toward a fully formed SNARE four-helix bundle.

Interestingly, the Cpx1(27-72) accessory helix exhibited clear steric clashes with the vesicle in all complexes. To avoid such clashes, the continuity between the central and accessory helices was broken in some cases, with the helix bending to one side or another (Figure 3G,I). In PC4, the entire helix changed orientation (Figure 3K) whereas in PC1, where the four-helix bundle is almost fully assembled, the helix was distorted into a snake shape (Figure 3E). The ‘struggle’ of the accessory helix to avoid bumps with the vesicle is particularly well illustrated by distinct bends of the Cpx1(27-72) helix occurring in PC3 during the simulations (Figure 3-figure supplement 2A-E). It is also noteworthy that Cpx1(27-72) remained bound to the SNAREs throughout the simulations due to interactions of the C-terminal end of the Cpx1(27-72) helix, particularly the Y70 aromatic ring, with a hydrophobic pocket of the SNARE complex, which persisted even when the overall direction of the helix changed in PC4 (Figure 3-figure supplement 2F). Overall, these observations provide a vivid visual illustration supporting the proposal that the complex-1 accessory helix inhibits fusion (Xue et al., 2007) because of its steric hindrance with the vesicle (Trimbuch et al., 2014).

A common feature of the four primed complexes at the end of the simulation was the arrangement of the synaptotagmin-1 C_2_B domain, which was initially placed between the SNARE four-helix bundle and the flat bilayer but in all primed complexes changed orientation, establishing extensive interactions between its polybasic face and the flat bilayer, and bringing the SNARE four-helix bundle close to the flat bilayer (Figure 4A-D). This arrangement dictates that the Cpx1(27-72) helix points towards the vesicle membrane, in agreement with the notion that binding of synaptotagmin-1 to the SNARE complex through the primary interface supports the inhibitory activity of synaptotagmin-1 and complexin-1 (Guan et al., 2017; Voleti et al., 2020). The synaptotagmin-1 C_2_A domain adopted distinct orientations in the different primed complexes, consistent with the fact that no stable Ca^2+^-independent interactions of this domain with membranes or the SNARE complex have been identified. The C_2_B domain consistently remained bound to the SNARE four-helix bundle via the primary interface in all four primed complexes throughout the simulation. The binding modes in the primed complexes resembled those observed in various crystal structures containing the primary interface (Zhou et al., 2015; Zhou et al., 2017), particularly in the so-called region I of this interface that includes Y338 among other side chains of C_2_B (e.g. Figure 4E). However, there were some differences in the other region of this interface (region II), which includes R281, R398 and R399 of the C_2_B domain. In the crystal structures, there was variability in the contacts made by these side chains and R398 did not interact with acidic residues or was at moderate proximity with E238 of syntaxin-1 (Figure 4-figure supplement 1A,B). However, the R398 side chain interacted with a negative pocket formed by E55, D58 and E62 of SNAP-25 in the four primed complexes of our simulation (Figure 4F, Figure 4-figure supplement 1C-E). The findings that an R398Q mutation impairs binding of the C_2_B domain to the SNARE complex in vitro (Voleti et al., 2020) and disrupts neurotransmitter release in neurons (Xue et al., 2008) support the relevance of the interactions of R398 uncovered by our simulation and suggest that crystal packing might have slightly distorted the binding mode, but it is also plausible that the binding mode is dynamic in this area.

**Figure 4.**
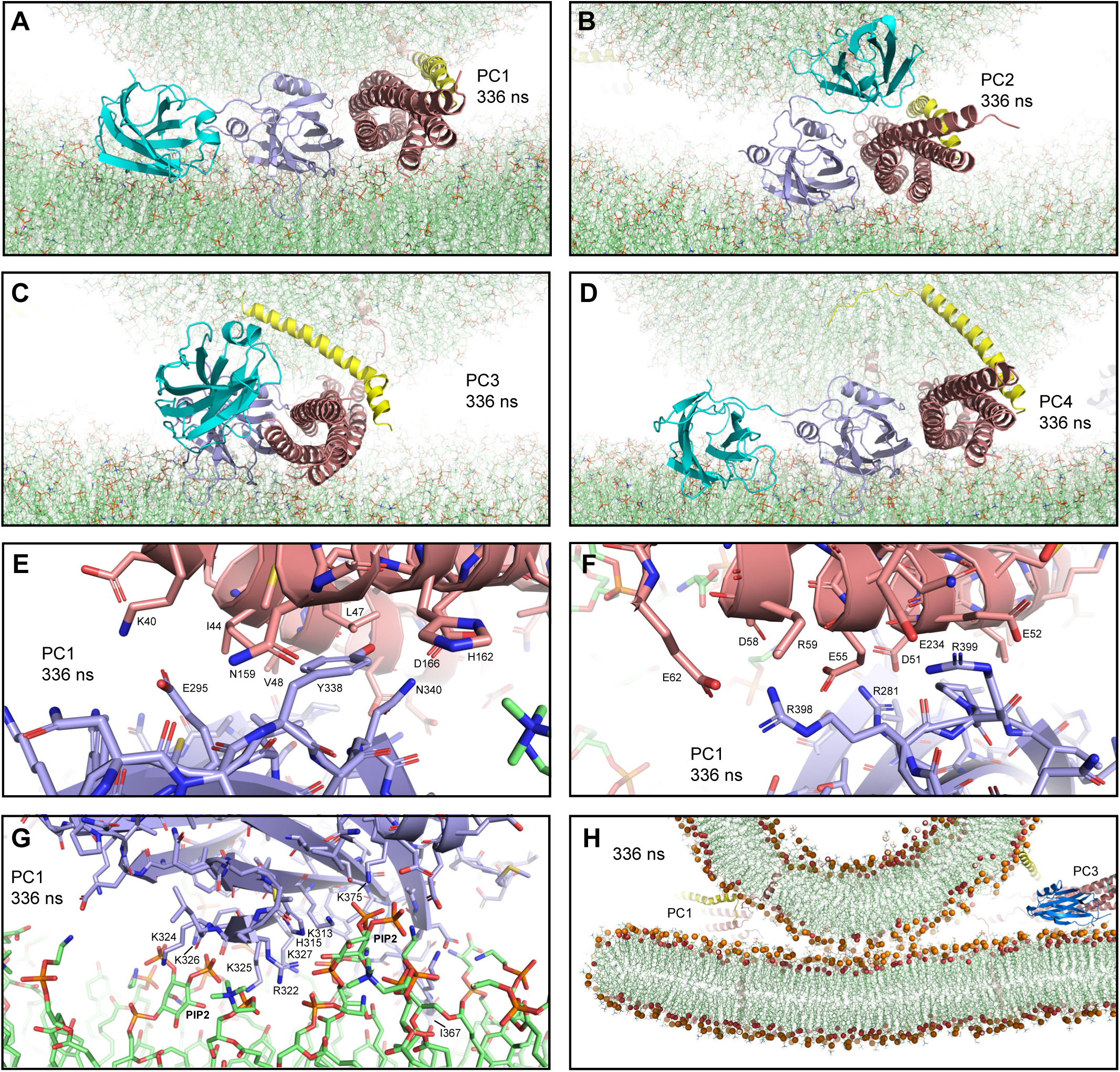
Additional views of the first MD simulation of primed complexes bridging a vesicle and a flat bilayer. (**A-D**) Close up views of the four primed complexes after 336 ns showing how the synaptotagmin-1 C_2_B domain binds to the SNARE complex through the primary interface and to the flat bilayer with the polybasic face, which dictates that the Cpx1(27-72) helix is oriented toward the vesicle and bends in different ways and directions to avoid steric clashes. This arrangement forces the SNARE four-helix bundle to be close to the flat bilayer. The primed complexes are illustrated by ribbon diagrams, with the SNAREs in salmon, Cpx1(27-72) in yellow and the synaptotagmin-1 C_2_AB fragment in cyan (C_2_A domain) and violet (C_2_B domain). The lipids are shown as thin stick models (carbon lime, oxygen red, nitrogen blue, phosphorous orange). (**E-F**) Two different close up views of the primary interface between the C_2_B domain and the SNARE complex in PC1 after 336 ns showing site I of the interface (**E**) or site II where R398,R399 of the C_2_B domain are located (**F**). The C_2_B domain and the SNARE complex are illustrated by ribbon diagrams and stick models with oxygen atoms in red, nitrogen atoms in blue, sulfur atoms in light orange and carbon atoms in violet [for the C_2_B domain] or salmon (for the SNAREs). The positions of selected side chains are indicated. (**G**) Close up view of the interaction of the C_2_B domain of PC1 with the flat bilayer after 336 ns. The positions of PIP_2_ headgroups, basic side chains involved in interactions with the lipids, and the hydrophobic side chain of I367 at the tip of a Ca^2+^-binding loop that inserts into the bilayer, are indicated. (**H**) Thin slice of the system showing a point-of-contact interface between the vesicle and the flat bilayer at 336 ns. Phosphorous atoms of phospholipids and the oxygen atoms of cholesterol molecules are shown as spheres to illustrate the approximate locations of lipid head groups. The positions of PC1 and PC3 are indicated.

In all four primed complexes, the extensive interactions of the C_2_B domain with the flat bilayer involved not only a polybasic sequence (residues 321-327) known to bind to PIP_2_ (Bai et al., 2004) but also other basic residues on this face of the β-sandwich that are also important for neurotransmitter release [e.g. K313, (Brewer et al., 2015)] (Figure 4G, Figure 4-figure supplement 1F-H)]. PIP_2_ molecules of the flat bilayer were often involved in these interactions. In addition, for all primed complexes, one of the C_2_B domain Ca^2+^-binding loops interacted extensively with the flat bilayer, inserting the hydrophobic residue at its tip (I367) into the acyl region. We also observed some interactions of the flat bilayer with basic residues of the SNARE four-helix bundle (e.g. R30,R31 from synaptobrevin and R176 from SNAP-25), which appeared to be favored because the clashes between the Cpx1(27-72) helix and the vesicle push the SNARE four-helix bundle and C_2_AB toward the flat bilayer. There was some variability in the four-helix bundle-flat bilayer interactions observed in the different primed complexes, but the overall arrangement of the C_2_B domain with respect to the flat bilayer and the SNARE four-helix bundle was very similar in all complexes, regardless of the orientation of the Cpx1(27-72) helix and the state of assembly of the SNARE four-helix bundle at the C-terminus (Figure 4A-D).

To further test the consistency of our results with respect to the configuration of the primed state, we built a similar system but using different configurations of the four initial primed complexes, which we generated by performing a restrained MD simulation analogous to the one that yielded the four initial primed complexes described above (Figure 3-figure supplement 1), but using a different random seed number and running the simulation for 50 ns instead of 1 ns. Moreover, we built a slightly larger square bilayer (35 x 35 nm^2^) to provide more space for protein-membrane interactions. The final system (Figure 4-figure supplement 2A,B) contained 5.9 million atoms and was used to run a production simulation of 310 ns. Figure 4-figure supplement 2C shows the final configuration. The behaviors of the primed complexes were similar to those of the previous simulation. The four initial four-helix bundles again had different levels of assembly at the C-terminus, with PC1 being the only one that was almost completely assembled, and there was not much progress toward full assembly in the other three complexes (Figure 4-figure supplement 2D-K). The Cpx1(27-72) helix again exhibited strong clashes with the vesicle and distinct ways to overcome such clashes, whereas the C_2_B domain changed orientation to establish extensive interactions with the flat bilayer while remaining bound to the SNARE four-helix bundle via the primary interface (Figure 4-figure supplements 2D-K, 3A-D). It is also noteworthy that in the two simulations of primed complexes, contacts between the vesicle and the plasma membrane were established at about 210-230 ns and the contacts increased gradually afterwards, but appeared to be leveling off at the end of the simulations (Figure 4-figure supplement 4), resulting in point-of-contact interfaces between the vesicle and the flat bilayer, without flattening of the vesicle (Figure 4H, Figure 4-figure supplement 3E).

Overall, the arrangements of the synaptotagmin-1 C_2_B domain with respect to the flat bilayer and the SNARE four-helix bundle in the eight primed complexes from the two simulations were very similar, and in all cases dictated that the Cpx1(27-72) helix was oriented toward the vesicle (Figure 4A-D, Figure 4-figure supplement 3A-D). The consistency of these results, together with the abundant data available on the functional importance of the C_2_B-membrane, C_2_B-SNARE and Cpx1(27-72)-SNARE interfaces present in these complexes [e.g. (Chen et al., 2002; Li et al., 2006; Zhou et al., 2015)] suggest that these complexes resemble those present in the primed state of synaptic vesicles.

### Potential effects of Ca^2+^ binding to synaptotagmin-1

Ca^2+^ binding to synaptotagmin-1 is believed to induce a tight, PIP_2_-dependent interaction of the C_2_B domain with the plasma membrane and dissociation from the SNARE complex to relieve the inhibition of release caused by synaptotagmin-1 and complexin-1 (Voleti et al., 2020). To examine whether we could observe the dissociation step and investigate how the system evolves afterwards through MD simulations, we generated a system analogous to that used for our first simulation of primed complexes, with the trans-SNARE complexes and their bound C_2_AB molecules in the same positions as in Figure 3-figure supplement 1 and the same vesicle, but with the large flat bilayer of 35 x 35 nm^2^ used for the second simulation of primed complexes to provide sufficient room for Ca^2+^-dependent binding to the C_2_ domains. We added five Ca^2+^ ions to the corresponding binding sites of C_2_A and C_2_B, and we removed the Cpx1(27-72) molecules to facilitate potential eventual fusion and to study at the same time how the system evolves without complexin-1. The final system (Figure 5A) contained 5.9 million atoms.

**Figure 5.**
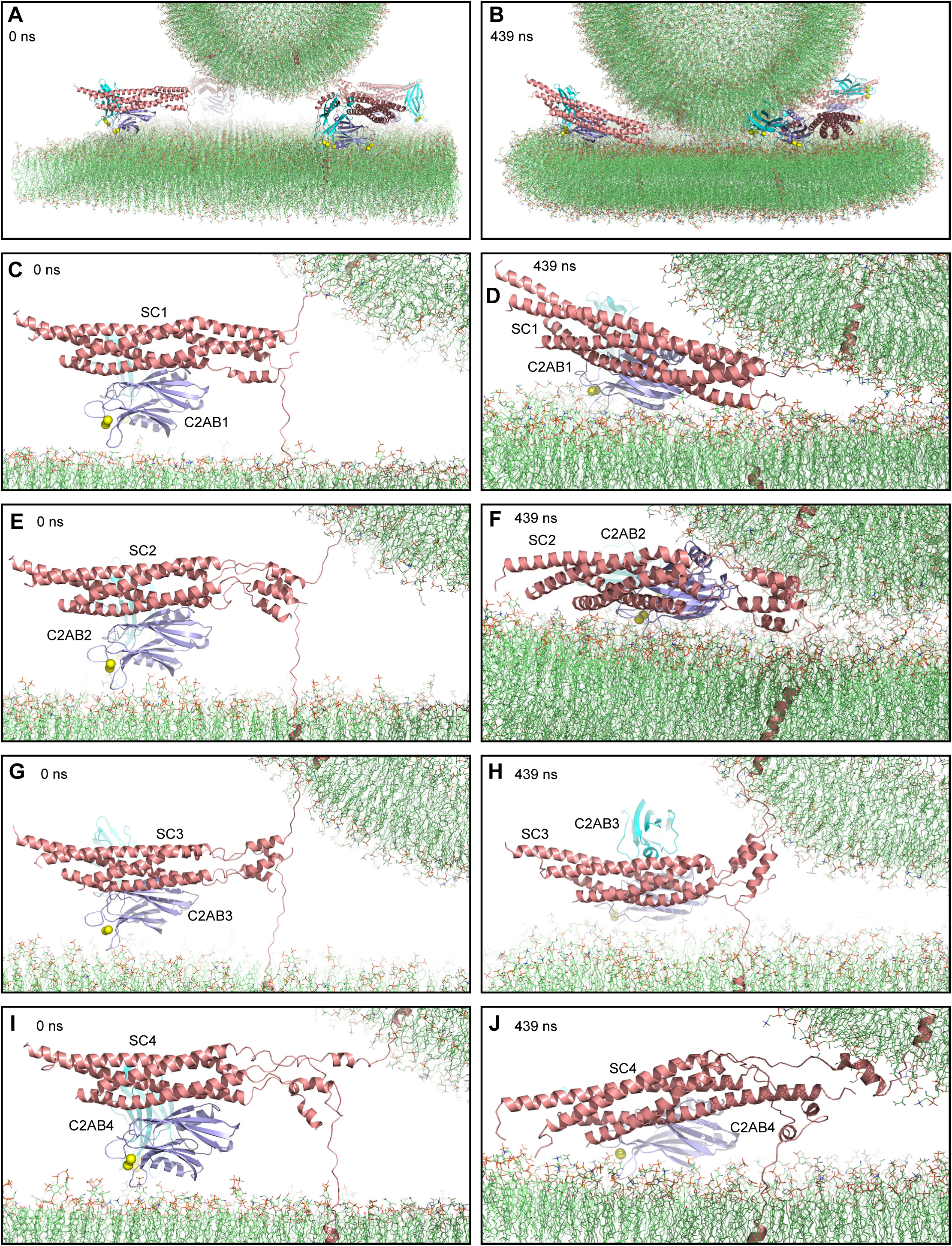
MD simulation of C_2_AB bound to Ca^2+^ and to trans-SNARE complexes bridging a vesicle and a flat bilayer. (**A**) Close up view of the four C_2_AB-SNARE complexes in the initial system. (**B**) Close up view of the system after a 439 ns MD simulation. (**C-J**) Close up views of the assemblies between C_2_AB molecules (named C2AB1-4) and SNARE complexes (SC1-SC4) in the initial configuration (**C,E,G,I**) and after the 439 ns MD simulation (**D,F,H,J**). In all panels, the SNAREs are represented by ribbon diagrams in salmon and the synaptotagmin-1 C_2_AB fragment by ribbon diagrams in cyan (C_2_A domain) and violet (C_2_B domain). Ca^2+^ ions are shown as yellow spheres. The lipids are shown as thin stick models (carbon lime, oxygen red, nitrogen blue, phosphorous orange).

We performed a production MD simulation of 439 ns, which led to the final configuration shown in Figure 5B. The C_2_AB-SNARE complexes generally behaved similarly to the primed complexes in the previous simulations, but with some differences. The SNARE four-helix bundle that was almost fully assembled in the starting configuration remained assembled almost completely (up to layer +7), whereas the other three complexes did not make much progress toward C-terminal assembly (Figure 5C-J). We note again that 439 ns are expected to provide ample time for helix formation and large conformational rearrangements, which is exemplified by the behavior of one of the SNARE four-helix bundles (SC4) during the simulation. Thus, the helix corresponding to the SNAP-25 C-terminal SNARE motif was almost fully formed after 5 ns, even though there was a substantial break in the helix in the beginning, and there were considerable structural changes at 75 ns, but only limited changes from 75 to 439 ns (Figure 5-figure supplement 1A-D). These findings again show that the constraints imposed by the system hinder fast assembly of the C-terminus of the SNARE four-helix bundle. Interestingly, SNARE four-helix bundles exhibited less interactions with the flat bilayer than in the simulations of primed complexes including Cpx1(27-72), consistent with the notion that the steric clashes of the complexin-1 accessory helix with the vesicle push the SNARE four-helix bundle toward the flat bilayer. As observed in previous simulations, the C_2_B domains of the four complexes established extensive interactions with the flat bilayer and remained bound to the SNARE complex through the primary interface (Figure 5D,F,H,J). We did observe that SC4 became detached from R398,R399 of the C_2_B domain early in the simulation and there were additional interactions in region I of the primary interface that remained at the end of the simulation (Figure 5-figure supplement 1E,F). However, it is unclear whether this change was caused by Ca^2+^ binding to the C_2_B domain. These findings suggest that dissociation of the C_2_B domain from the SNAREs requires longer time scales and may be a rate limiting step in release, which is supported by the finding that an E295A/Y338W in the C_2_B domain primary interface enhances SNARE complex binding (Voleti et al., 2020) but disrupts Ca^2+^-evoked neurotransmitter release (Zhou et al., 2015). In this simulation, the vesicle came into contact with the flat bilayer at about 400 ns and the number contacts increased gradually afterwards but without reaching a plateau at the end (Figure 5-figure supplement 2).

Ca^2+^-dependent interactions of both synaptotagmin-1 C_2_ domains with lipids are crucial for neurotransmitter release (Fernandez-Chacon et al., 2001; Rhee et al., 2005) and likely cooperate with the SNAREs in accelerating membrane fusion by a mechanism that is currently unknown. This function might be performed by the same synaptotagmin-1 molecules that dissociate from the SNARE complex or, perhaps more likely, by other molecules that were previously located near the vesicle-plasma membrane interface, poised to assist in membrane fusion. To explore these ideas through MD simulation, we generated a system that was identical to the starting point of the first simulation of primed complexes (Figure 3A), but removing the Cpx1(27-72) molecules and moving the C_2_AB molecules to arbitrary locations between the SNARE complexes, near the vesicle and the flat bilayer. Five Ca^2+^ ions were added to the corresponding binding sites of each C_2_AB. The system (Fig. 6A) contained 4.9 million atoms. After equilibration, we performed a 668 ns production MD simulation.

**Figure 6.**
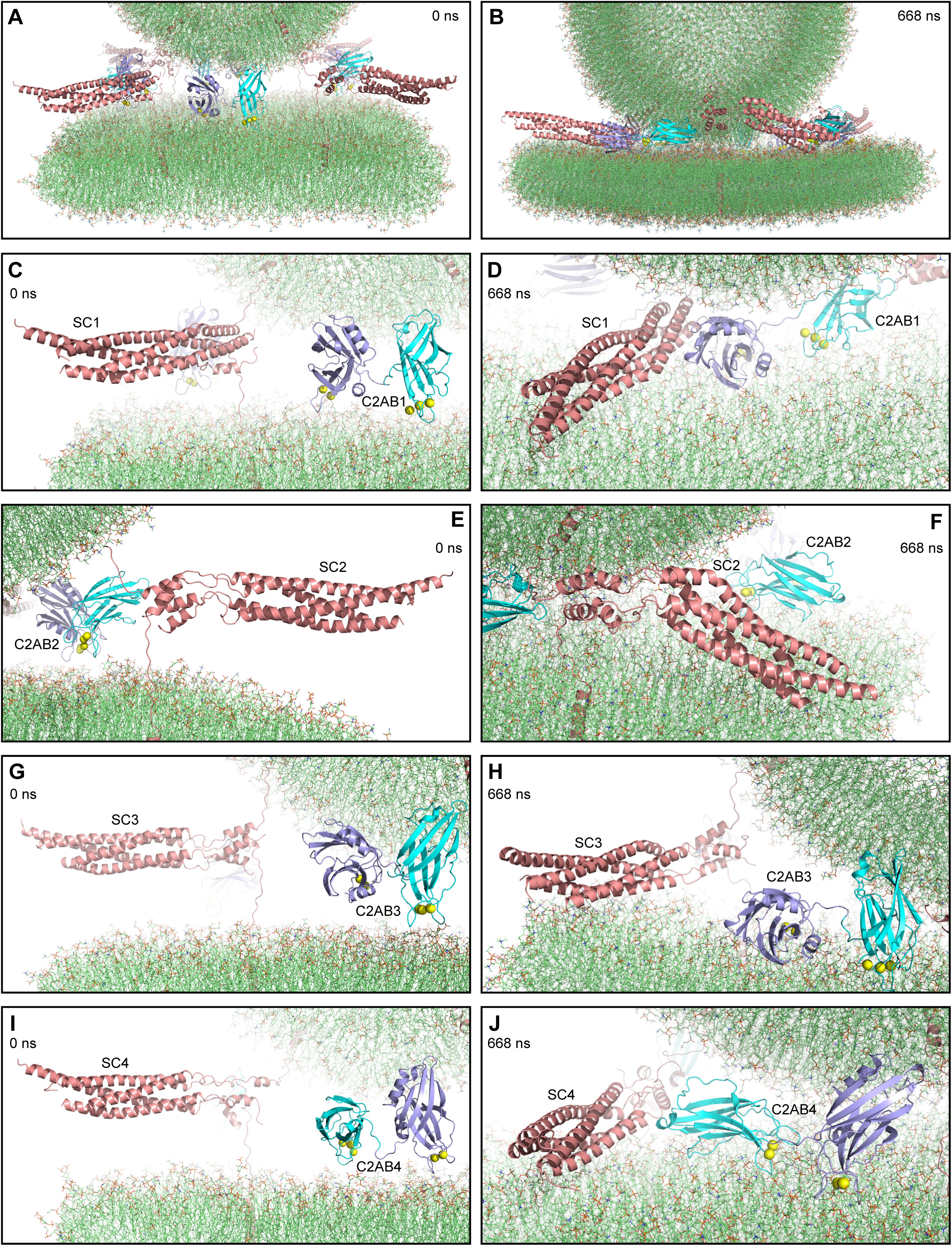
MD simulation of Ca^2+^-bound C_2_AB dissociated from trans-SNARE complexes bridging a vesicle and a flat bilayer. (**A**) Close up view of the four C_2_AB molecules and SNARE complexes in the initial system. (**B**) Close up view of the system after a 668 ns MD simulation. (**C-J**) Close up views of the individual C_2_AB molecules (named C2AB1-4) and SNARE complexes (SC1-SC4) in the initial configuration (**C,E,G,I**) and after the 668 ns MD simulation (**D,F,H,J**). In all panels, the SNAREs are represented by ribbon diagrams in salmon and the synaptotagmin-1 C_2_AB fragment by ribbon diagrams in cyan (C_2_A domain) and violet (C_2_B domain). Ca^2+^ ions are shown as yellow spheres. The lipids are shown as thin stick models (carbon lime, oxygen red, nitrogen blue, phosphorous orange).

The SNARE complexes behaved similarly as in other simulations (Figure 6C-J, Figure 6-figure supplement 1). The distinct C_2_AB molecules exhibited different degrees of changes with respect to their initial states, as well as different orientations of their C_2_ domains with respect to each other and to the membranes. Interestingly, the C_2_B domain of one of the C_2_AB molecules (C_2_AB1) bound to the C-terminal half of the SC1 four-helix bundle through a surface similar to the primary interface and including interactions of R398 with acidic residues of synaptobrevin (Figure 6-figure supplement 1A,C). This interaction is interesting because it orients the Ca^2+^-binding loops of the C_2_B domain toward the vesicle-flat bilayer interface, close to the area where the juxtamembrane sequence of synaptobrevin containing three aromatic residues inserts into the vesicle, and the Ca^2+^-binding loops of the C_2_A domain are also oriented toward this interface (Figure 6D, Figure 6-figure supplement 1A,B). This arrangement could facilitate cooperation between the juxtamembrane region of synaptobrevin and the Ca^2+^-binding loops of the synaptotagmin-1 C_2_ domains in perturbing the lipid bilayers to initiate membrane fusion, as proposed in a previous model (Rizo and Xu, 2015). We note that this interaction was not observed in NMR studies in solution (Brewer et al., 2015; Voleti et al., 2020) and is incompatible with complexin-1 binding because of steric hindrance, but this binding mode might be facilitated by interactions with the membranes, and some data suggested that C_2_AB competes with complexin-1 for binding to membrane-anchored SNARE complexes by binding to their C-termini (Dai et al., 2007).

C_2_AB2, C_2_AB3 and C_2_AB4 had limited contacts with the adjacent SNARE complexes, but all interacted extensively with the flat bilayer through their Ca^2+^-binding loops, as did C_2_AB1 (Figure 6C-J, Figure 6-figure supplement 1). In addition, the polybasic region of three of the C_2_B domains established extensive contacts with the flat bilayer that led to slanted or parallel orientations with respect to the flat bilayer, but the C_2_B domain of the C_2_AB4 molecule adopted a more perpendicular orientation (Figure 6J) that resembles the orientation predicted according to the distinct effects of mutations in the polybasic region on binding to PIP_2_-containing membranes (Voleti et al., 2020). This orientation allows interactions between the vesicle and the region of the C_2_B domain containing R398,R399, which is opposite to the Ca^2+^-binding loops. The C_2_A domain of the C_2_AB3 molecule also bridges the vesicle and the flat bilayer in a similar orientation (Figure 6H). Both C_2_ domains of the C_2_AB1 molecule interacted with the vesicle through other areas of their β-sandwich structures. Because several of the interactions involve areas of the C_2_ domains outside of the Ca^2+^-binding loops, and because such interactions might be favored by the localization of synaptotagmin-1, it is plausible that synaptotagmin-1 molecules that do not interact with the SNARE complex may be bound to the vesicle and the plasma membrane before Ca^2+^ influx. It is tempting to speculate that, through such interactions, synaptotagmin-1 may act as a wedge that prevents extension of the membrane-membrane interface before Ca^2+^ influx and at the same time is ready to quickly facilitate fusion when Ca^2+^ binding induces insertion of the Ca^2+^-binding loops into the bilayers, causing perturbations that add to those caused by the SNARE complex to induce fast membrane fusion. Indeed, although the vesicle and the flat bilayer came into contact at about 360 ns and the number of contacts increased gradually afterwards, the increase was gradual (Figure 6-figure supplement 2) and a point-of-contact interface between the vesicle and the flat bilayer was still observed at the end of the simulation (Figure 6-figure supplement 1G). However, longer simulations will be required to confirm that synaptotagmin-1 prevents the formation of an extended vesicle-flat bilayer interface.

## Discussion

The high speed of Ca^2+^-evoked neurotransmitter release is possible because synaptic vesicles are primed to a metastable state that is ready for very fast fusion with the plasma membrane upon Ca^2+^ influx. Priming involves the formation of trans-SNARE complexes between the vesicle and plasma membranes, which is orchestrated by Munc18-1 and Munc13-1 (Prinslow et al., 2019). Synaptotagmin-1 and complexin-1 are believed to be bound to trans-SNARE complexes in the primed state, keeping the system ready for fast fusion but preventing premature fusion before Ca^2+^ influx (Voleti et al., 2020). Although X-ray crystallography has provided key structural information on the protein-protein interactions underlying this state (Chen et al., 2002; Zhou et al., 2015), the configuration of this state with regard to the vesicle and plasma membranes is unclear, and the steps that lead to membrane fusion upon Ca^2+^ influx remain highly enigmatic. Our MD simulations do not provide definitive conclusions to address these questions, but they help visualize potential configurations of the system in different stages leading to fusion and yield multiple clues on structural and mechanistic factors that may be determinant for fast membrane fusion. In particular, the simulations suggest that trans-SNARE complexes alone strongly draw two membranes together but their action may lead to formation of extended vesicle-plasma membrane interfaces that are expected to fuse slowly. Furthermore, our results suggest that trans-SNARE complexes are close to fully zippered in the primed state and form macromolecular assemblies with synaptotagmin-1 bound on one side and complexin-1 bound on the other side, in a spring-loaded arrangement where interactions of the synaptotagmin-1 C_2_B domain with the plasma membrane dictate that the complexin-1 helix bumps with the vesicle, hindering progression of the system toward fusion.

Our results need to be interpreted with caution because of the limited simulation times, as well as uncertainties about the initial configurations and the accuracy the force field used. Nevertheless, many of the features that we observed make sense from structural and energetic points of view, and are consistent with abundant experimental data available on this system. The short restrained simulation used initially to generate the trans-SNARE complexes led to unwinding of the helical conformations of the juxtamembrane linkers of synaptobrevin and syntaxin-1 while the SNARE four-helix bundle remained assembled (Figure 1-figure supplement 1A), which is consistent with the high stability of the SNARE complex (Gao et al., 2012). The juxtamembrane linkers remained unstructured in all our simulations but established extensive contacts with the membranes due to the abundance of basic residues in their sequences, and of aromatic residues in the case of the synaptobrevin linker. These linker-lipid interactions likely contributed to quickly (within 270 ns) bring the two flat bilayers together such that the lipids packed against the SNARE four-helix bundles (Figure 1C,D). However, we did not observe the initiation of fusion in the 750 ns that we simulated. We again did not observe fusion in a simulation of four trans-SNARE complexes bridging a vesicle and a flat bilayer performed at 310 K (520 ns) or in an ensuing simulation performed at 325 K (454 ns), but the complexes did draw the two membranes closely together early in the 310 K simulation, leading to flattening of the vesicle and formation of an extended vesicle-flat bilayer interface (Figure 2J). These results are consistent with the expectation that SNARE complexes act as powerful engines to draw two membranes together and suggest that this action depends not only on the high stability of the SNARE four-helix bundle but also on interactions of the juxtamembrane linkers with the lipids. However, our data suggest that the SNAREs alone are not able to induce fusion in the low microsecond time scale, at least in part because they have a tendency to cause formation of extended membrane-membrane interfaces. This conclusion is supported by the finding that, although such interfaces have been observed experimentally, they evolve to fusion in slow time scales (seconds-minutes) (Hernandez et al., 2012; Witkowska et al., 2021).

These observations suggest that other proteins involved in Ca^2+^-evoked release may help to prevent the formation of extended vesicle-plasma membrane interfaces, which is also supported by cryo-EM images of fusion reactions using reconstituted liposomes (Gipson et al., 2017). Hence, setting up a system that resembles the primed state of synaptic vesicles may be critical to recapitulate the steps that lead to fast Ca^2+^-triggered synaptic vesicle fusion in microsecond-scale MD simulations. The nature of the primed state is unclear, but strong evidence indicates that this state includes complexin-1 bound to one side of the SNARE complex as revealed by a crystal structure (Chen et al., 2002; Xue et al., 2007), and synaptotagmin-1 bound to the opposite side of the SNARE complex via the primary interface of the C_2_B domain (Zhou et al., 2015). A model of the primed state based on these crystal structures and the known tendency of the C_2_B domain to bind to PIP_2_-containing membranes (Bai et al., 2004) envisioned that the SNARE complex is ‘sandwiched’ between the C_2_B domain and complexin-1, which hinders C-terminal zippering (Voleti et al., 2020). This model was used to build the system for our two MD simulations of the primed state (Figure 3A,B, Figure 4-Figuresupplement 2A,B), which together included eight different states of C-terminal assembly of the SNARE four-helix bundle. Common features of the eight primed complexes at the late stages of the simulations included the extensive interactions of the C_2_B domain polybasic face with the flat bilayer, the persistence of the binding of the C_2_B domain to the SNAREs through the primary interface and the reorientation of the system such that the SNARE four-helix bundle came near the flat bilayer, with the C_2_B domain on the side rather than between the bundle and the flat bilayer, and with the helix of Cpx1(27-72) oriented toward the vesicle membrane (Figures 3, 4, Figure 4-figure supplement 2). The consistency of these features regardless of the state of C-terminal assembly of the SNARE four-helix bundle suggests that they are characteristics of the primed state of synaptic vesicles and that at least some of the configurations of the primed complexes visited in the simulations resemble those present in the primed state.

We did observe some variability in the binding mode between the C_2_B domain and the SNARE complex, as in some of the primed complexes the R398 side chain exhibited interactions with an acidic pocket formed by E55, D58 and E62 of SNAP-25 that were not observed by crystallography (e.g. Figure 4F). However, these represent small differences that did not have substantial consequences for the relative orientation of the C_2_B domain and the SNARE complex. Large variations were observed in the Cpx1(27-72) helix, which bent at different places or in different directions to avoid steric clashes with the vesicle. These differences may be due to random motions but they may also have arisen in part from the different states of assembly at the C-terminus of the four-helix bundle. It is quite remarkable that, despite the steric clashes, the Cpx1(27-72) helix did not become unstructured throughout both simulations for any of the complexes. These observations correlate with the finding that the sequence of the accessory helix of complexin-1 has a high intrinsic tendency to form α-helical structure, which nucleates helical structure in the region of the central helix (Pabst et al., 2000; Radoff et al., 2014). Overall, the configurations of the eight primed complexes provide a clear illustration of the notion that the complexin-1 accessory helix can have steric clashes with the vesicle, which was proposed to underlie the inhibition of neurotransmitter release by complexins (Trimbuch et al., 2014). Recent cross-linking experiments suggested that the inhibition arises because interactions of the complexin-1 accessory helix with synaptobrevin and SNAP-25 hinder C-terminal zippering (Malsam et al., 2020). However, it is unlikely that specific protein-protein interactions underlie the inhibitory function, as replacement of the complexin accessory helix with a completely unrelated sequence that has a high tendency to form α-helical conformation did not impair the inhibition of release by complexin in *C. elegans*, and helix propensity appears to be the key determinant for the inhibitory function (Radoff et al., 2014). These findings are fully consistent with the steric hindrance model supported by previous functional data (Trimbuch et al., 2014) and our MD simulations.

The extent of C-terminal assembly of the SNARE four-helix bundle was different in the eight initial complexes of the two simulations of the primed state generated to provide different starting points and explore whether C-terminal assembly could occur in the time scale of the simulations. Interestingly, the four-helix bundle was disrupted in the C-terminal half while the very C-terminus remained assembled in several of the initial complexes and this feature was maintained during the simulations (Figure 3, Figure 4-figure supplement 2). This finding suggests that the very C-terminus of the four-helix bundle is particularly stable. The disrupted region of the four-helix bundle did not progress much toward assembly in any of these complexes or the ones where the bundle was even more disrupted. Since helices can readily form and large conformational changes can occur in unstructured regions in the 100 ns time scale (Munoz and Cerminara, 2016), these observations indicate that completion of C-terminal assembly did not occur in these complexes because of the constraints imposed by the system. Note also that one complex of each simulation was almost completely assembled from the beginning to the end even though the Cpx1(27-72) helix was bumping into the vesicle as the two membranes came closer (Figure 3D,E, Figure 4-figure supplement 2D,E). However, as illustrated in the close-up views of Figure 7A-C, the vesicle and the flat bilayer were not as close to each other at the end of the two simulations of primed complexes as they were in the simulation of a vesicle and a flat bilayer with four trans-SNARE complexes alone after 280 ns at 310 K, when the extended vesicle-flat bilayer interface had already formed (Figure 2J).

**Figure 7.**
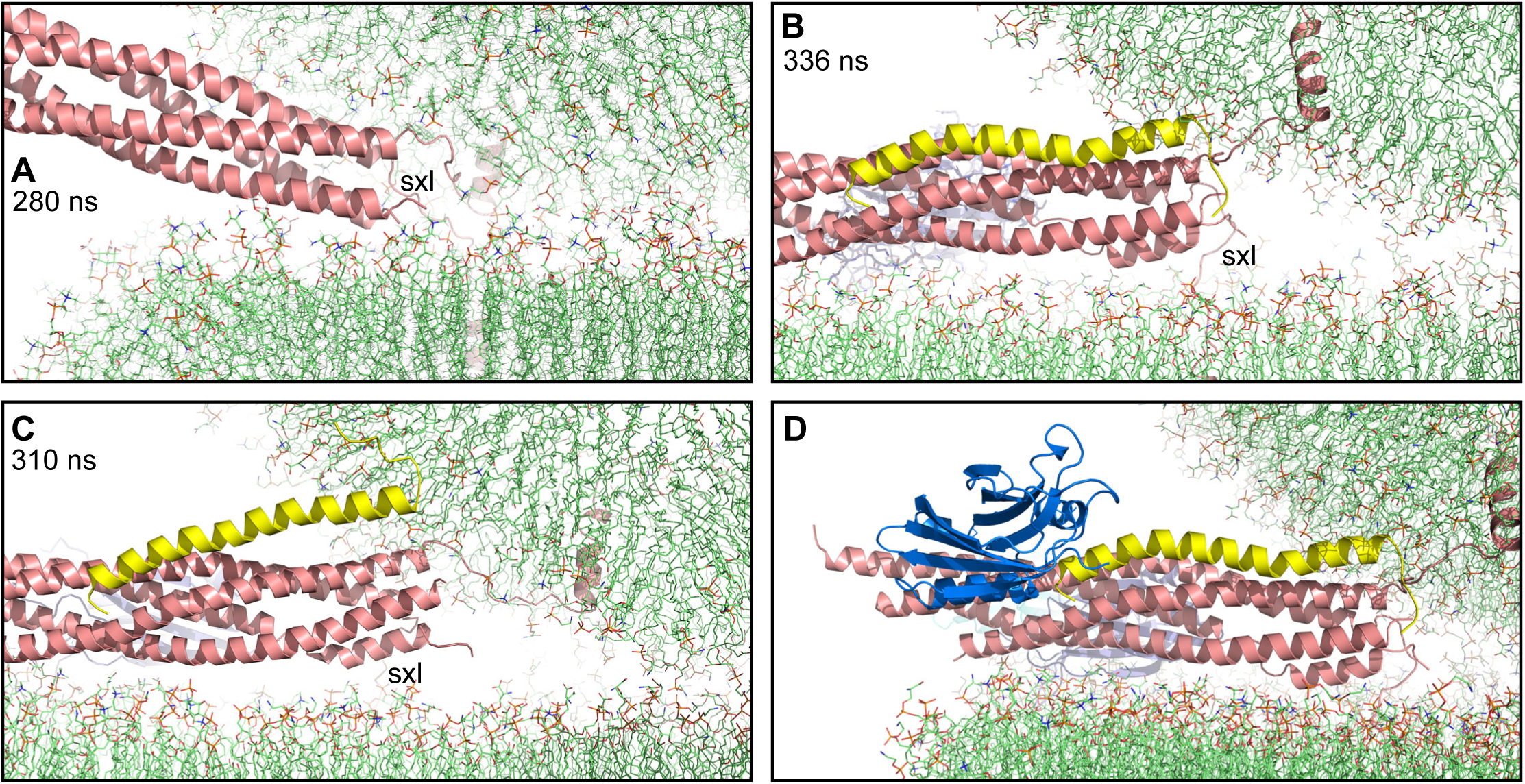
Complexin-1 may hinder the final action of trans-SNARE complexes to bring membranes together. (**A-C**) Close up views of one of the SNARE complexes bridging a vesicle and a flat bilayer after simulation for 280 ns at 310 K (**A**; shown in Figure 2D after 520 ns at 310 K and 454 ns at 325 K), of PC1 in the first MD simulation of primed complexes after 336 ns (**B**, also shown in Figure 3E) and of PC1 in the second MD simulation of primed complexes after 310 ns (**C**, also shown in Figure 4-figure supplement 2E). The complexes are illustrated by ribbon diagrams, with the SNAREs in salmon, Cpx1(27-72) in yellow and the synaptotagmin-1 C_2_AB fragment in cyan (C_2_A domain) and violet (C_2_B domain). The positions of the syntaxin-1 juxtamembrane linkers (sxl) are indicated. The comparison shows how the SNARE complex with the fully assembled four-helix bundle in (**A**) drew the two membranes closer than the two primed complexes of (**B,C**). (**D**) Close up view illustrating the position that would be occupied by the C_2_B domain of a second C_2_AB fragment (blue; only C_2_B domain shown) bound to the primed complex of (**B**) through the tripartite interface.

It is tempting to speculate from these observations that, to maximize the speed of release, SNARE complexes are almost fully zippered in the primed state of synaptic vesicles before Ca^2+^ influx, as in the PC1 primed complexes at the end of the simulations (Figure 7B,C), and that complexin-1 stabilizes this state but hinders the ‘final pull’ of the SNARE four-helix bundle to bring the membranes together, preventing the formation of extended interfaces and at the same time keeping the system ready for fast fusion upon Ca^2+^ influx. In this spring-loaded model, binding of the synaptotagmin-1 C_2_B domain to the SNARE complex via the primary interface contributes to form this state through its interactions with the plasma membrane and the SNAREs, which dictate the orientation of complexin-1 toward the vesicle and hinder rotations of the four-helix bundle that could eliminate the steric clashes between complexin-1 and the vesicle. Upon Ca^2+^ influx, dissociation of the Ca^2+^-bound C_2_B domain from the SNARE complex (Voleti et al., 2020) allows such rotations and both C_2_ domains can cooperate with the SNAREs in triggering fusion by a mechanism that remains unclear (see below). Hence, this model can explain the finding that complexins are required for the dominant negative effect of mutations in the Ca^2+^-binding site of the synaptotagmin-1 C_2_B domain (Zhou et al., 2017). We propose that this model corresponds to the tight primed state of synaptic vesicles that has been proposed to be in dynamic equilibrium with a loose primed state where SNARE complexes are less zippered at the C-terminus (Neher and Brose, 2018).

Clearly, this model will need to be tested with further MD simulations and experimentation, and there are other aspects of the primed state that remain to be resolved. For example, the C_2_B domain can also bind to the SNARE complex through a so-called tripartite interface whereby an α-helix of the C_2_B domain is adjacent to the complexin-1 helix (Zhou et al., 2017). Although binding through this interface was not detected by NMR spectroscopy at 85 μM concentration and hence appears to be very weak (Voleti et al., 2020), the affinity could be enhanced by interactions of synaptotagmin-1 with the vesicle (Brunger et al., 2018). Adding a C_2_B domain to our model of the primed state, bound to the SNARE complex via the tripartite interface, suggests that this C_2_B domain would be too far to interact with the vesicle (Figure 7D). However, the C_2_A domain (not shown) might bind to the vesicle. Moreover, it is also plausible that the vesicle and plasma membranes are not in contact, in contrast to our model, and the SNARE complexes are closer to the center of the vesicle-plasma membrane interface, which might place the C_2_B domain sufficiently close to bind to the vesicle. Note also that, although complexin-1 has very high affinity for the SNARE complex (Pabst et al., 2002), the steric clashes with the vesicle likely decrease the effective affinity but. In turn, the affinity could be increased by binding of the C_2_B domain to the tripartite interface. Hence, the tripartite complex also provides an explanation for the requirement of complexin for the dominant negative effects of C_2_B Ca^2+^-binding mutants (Zhou et al., 2017). An additional model that has been proposed for an inhibitory role of synaptotagmin-1 is based on the observations that synaptotagmin-1 forms oligomeric rings that are disrupted upon Ca^2+^ binding and that a mutation that impairs oligomerization (F349A) enhances spontaneous neurotransmitter release (Tagliatti et al., 2020). However, it is unclear whether there are sufficient synaptotagmin-1 molecules in synaptic vesicles (Takamori et al., 2006) to oligomerize and bind to both sites of the SNARE complex. Moreover, F349 is integral part of the tripartite interface and, therefore, the phenotype caused by the F349A mutation might arise from disruption of the tripartite complex. Hence, the physiological relevance of the tripartite complex and of synaptotagmin-1 oligomerization need to be tested further.

The events that lead to rapid fusion upon Ca^2+^ influx also remain unclear, but our simulations provide clues that may be key to understand these steps. In the 439 ns simulation with Ca^2+^-bound C_2_AB and the C_2_B domains bound to the SNARE complexes we did not observed Ca^2+^-induced dissociation from the SNARE complex (Figure 5), suggesting that such dissociation may be a rate limiting step in Ca^2+^-evoked release. This proposal is consistent with the finding that the E295A/Y338W mutation in the C_2_B domain enhances SNARE complex binding (Voleti et al., 2020) but disrupts Ca^2+^-evoked release (Zhou et al., 2015). Since the Ca^2+^-binding loops of the C_2_B domain bound to the SNARE complex are pointing away from the membrane-membrane interface where fusion occurs, it seems likely that the active role in membrane fusion to trigger neurotransmitter release is played by other synaptotagmin-1 molecules that are not bound to the SNARE complex and are close to the membrane-membrane interface due to their affinity for the PIP_2_-containing plasma membrane and their anchoring on the vesicle. The 668 ns MD simulation performed to test this notion did not reveal substantial membrane perturbations that could lead to fusion, but showed two interesting features that might be important for fusion. On the one hand, one of the C_2_AB molecules bound to the C-terminus of the SNARE complex (Figure 6D), an interaction that would require displacement of the complexin-1 accessory helix and might allow cooperation between the SNARE juxtamembrane regions and the C_2_ domain Ca^2+^-binding loops in perturbing the lipid bilayers to initiate fusion. Although there is some experimental evidence for this type of interaction in vitro (Dai et al., 2007), its functional relevance needs to be investigated. On the other hand, two of the C_2_AB molecules were found to bridge the vesicle and the flat bilayer through either the C_2_A domain or the C_2_B domain (Figure 6H,J). Through such bridging, synaptotagmin-1 could act as a wedge that prevents extension of the vesicle-plasma membrane interface while the SNARE complexes draw the two membrane together strongly, and the torque caused by these forces may help to induce local membrane bending to initiate fusion.

While geometrical arguments and the vesicle-flat bilayer contacts observed in our simulations (Figure 2-figure supplement 1D, Figure 4-figure supplement 4 and Figure 6-figure supplement 2) support the notion that bilayer adhesion leading to extended membrane-membrane interfaces may be prevented by interactions of the synaptotagmin-1 C_2_ domains with the membranes and/or by the primed assemblies formed by synaptotagmin-1, complexin and trans-SNARE complexes, more systematic analyses with longer simulations, together with experimental studies, will be required to test this proposal. Moreover, there are undoubtedly other possibilities for the mechanism of fast Ca^2+^-triggered membrane fusion beyond those described above. For instance, part of the syntaxin-1 linker is not interacting with the membranes in the states visited in our simulations (e.g. Figure 6-figure supplement 1B, Figure 7A-C) and rotation of the four-helix bundle to favor such interactions might be important to initiate fusion. It is also plausible that the SNAREs alone can induce fast fusion in configurations that we did not study, for instance if the SNARE TM regions were located closer to the center of the vesicle-flat bilayer interface. We hope that the work presented here and the coordinates of the systems that we have generated will provide a valuable resource to test these and other ideas on the mechanism of Ca^2+^-triggered synaptic vesicle fusion using MD simulations and experimentation.

## Methods

### High performance computing

Most high performance computing, including all production MD simulations, were performed using Gromacs (Pronk et al., 2013; Van Der Spoel et al., 2005) with the CHARMM36 force field (Huang et al., 2017; Klauda et al., 2010; Lee et al., 2019; Wu et al., 2014a; Wu et al., 2014b) on Frontera at TACC. Some of the initial setup tests, solvation, ion addition, minimizations and equilibration steps were performed using Gromacs with the CHARMM36 force field at the BioHPC supercomputing facility of UT Southwestern, or on Lonestar5 or Stampede2 at TACC. System visualization and manual manipulation were performed with Pymol (Schrödinger, LLC).

### System setup

All systems were built by manually combining coordinates of the protein components with coordinates of the membranes, solvating the system with explicit water molecules (TIP3P model) and adding potassium and chloride ions as needed to reach a concentration of 145 mM and make the system neutral. Flat lipid bilayers were built with the Membrane Builder module (Jo et al., 2007; Jo et al., 2009) in the CHARMM-GUI (Jo et al., 2008) website (https://charmm-gui.org/), providing the coordinates of the TM region of synaptobrevin or syntaxin-1 in their desired positions as input. The bilayers contained mixtures of cholesterol (CHL1), 16:0-18:1 phosphatidylcholine (POPC), 18:0-22:6 phosphatidyltethanolamine (SDPE), 18:0-22:4 phosphatidyltethanolamine (SAPE), 18:0-18:1 phosphatidylserine (SOPS), 18:0-22:6 phosphatidylserine (SDPS), 18:0-20:4 phosphatidylinositol 4,5-bisphosphate (SAPI2D) and/or 18:0-20:4 glycerol (SAGL). Table 1 list the number of atoms and lipid compositions of the membranes; each entry corresponds to a system denoted by the abbreviations described below, which were used as roots for the filenames of the corresponding simulations.

*Four trans-SNARE complexes bridging two flat bilayers* (qscff system). The starting point to generate a trans-SNARE complex was the crystal structure of the neuronal SNARE complex that included the TM regions of synaptobrevin and syntaxin-1 (Stein et al., 2009). Two residues at the C-terminus of syntaxin-1 (residues 287-288) were added in Pymol, and four residues of the C-terminus of SNAP-25 (residues 201-204) were added manually based on a crystal structure of soluble SNARE complex (PDB accession code 1NS7). The resulting complex included residues 30-116 of synaptobrevin, residues 189-288 of syntaxin-1, and residues 8-82 and 141-204 of SNAP-25. To move the TM regions of synaptobrevin and syntaxin-1 to designed positions where they were later inserted into the flat lipid bilayers (Figure 1-figure supplement 1A), the cis-SNARE complex was solvated with explicit water molecules and energy minimized. Then a 1 ns production MD simulation at 310 K was performed imposing position restraints to keep of all heavy atoms of the N-terminal half of the SNARE complex, up to the polar layer (residues 30-56 of synaptobrevin, residues 189-226 of syntaxin-1, and residues 8-53 and 141-174 of SNAP-25), in their original coordinates (force constant 1,000 kJ/mol/nm^2^), and to force the backbone atoms of the TM regions (residues 95-116 of synaptobrevin and residues 266-288 of syntaxin-1) to move to the designed positions (force constant 300 kJ/mol/nm^2^). Four copies of the final structure were rotated and translated to desired positions (Figure 1-figure supplement 1B), and merged with two square flat bilayers of 26 x 26 nm^2^ each.

*Four trans-SNARE complexes bridging two flat bilayers including four Ca^2+^-bound C_2_AB molecules* (Sqscff system). To generate C_2_AB molecules for this simulation, the C_2_AB molecule from a complex with the SNAREs (PDB accession number 5CCH) (Zhou et al., 2015) was used as a starting point. Ca^2+^ ions were added at the corresponding sites as observed in the solution NMR structures of the C_2_A and C_2_B domains (PDB accession codes 1BYN and 1K5W) (Fernandez et al., 2001; Shao et al., 1998). After solvation with explicit water molecules and energy minimization, a 10 ns production MD simulation was carried out using position restraints to keep the initial coordinates of the C_2_B domain and move the C_2_A domain to a designed location so that the Ca^2+^-binding loops of both C_2_ domains point in similar directions (Figure 1-figure supplement 1E) and hence can bind in a Ca^2+^-dependent manner to the same membrane (force constant 1,000 kJ/mol/nm^2^). The final C_2_AB structure was energy minimized and four copies of it were incorporated into the system containing four trans-SNARE complexes bridging two flat bilayers (qscff), interspersed between the SNARE complexes.

*Four trans-SNARE complexes bridging a vesicle and a flat bilayer* (qscv system). The vesicle was built by adaptation of the scripts for building coarse-grained vesicle systems from CHARMM-GUI Martini Maker (Qi et al., 2015). The radius of the vesicle was set to 11 nm and the number of lipids in the inner and outer layer of the vesicle, given the specific vesicle radius and lipid ratio in each layer, were calculated using the same scheme as in Martini Maker (Table 1). As the final system was too large for long-time equilibration of the lipids in the inner and outer layer using water pores along the x, y and z axis, no water pore was created in the vesicle (i.e., water pore radius was set to 0 nm). The trans-SNARE complex built for the qscff system was used, after solvation, minimization and equilibration, as a starting point for a 2 ns production MD simulation imposing position restraints to keep all heavy atoms of the N-terminal half of the SNARE complex in their original coordinates and to force the backbone atoms of the TM regions of synaptobrevin and syntaxin-1 to move to designed positions (force constant 1,000 kJ/mol/nm^2^). Four copies of the final structure were rotated and translated to designed positions (Figure 2-figure supplement 1A), and merged with the vesicle and a square flat bilayer of 26 x 26 nm^2^ each. After solvation and equilibration, we performed a 7 ns production MD simulation and observed the appearance of holes in the vesicle that arose because the lipid density was not optimal. The holes were filled manually with lipid patches from the original vesicle and a 5 ns production MD simulation was carried out with position restraints to keep the SNAREs in their initial locations. New holes appeared and were filled manually again. After another 10 ns production MD simulation with position restraints on the SNARE coordinates, no additional holes appeared. A final 80 ns production MD simulation with position restraints on the SNAREs was performed to equilibrate the vesicle lipids, which yielded the initial equilibrated system (Figure 2-figure supplement 1B,C) that was used to initiate an unrestrained production MD simulation.

*First simulation of primed complexes bridging a vesicle and a flat bilayer* (prsg system). The Cpx1(27-72) fragment was built starting from the crystal structure of Cpx1(26-83) bound to the SNARE complex (PDB accession code 1KIL) (Chen et al., 2002), which contained electron density for residues 32-73 of complexin-1. Five additional N-terminal residues in a random conformation were added in Pymol to yield Cpx1(27-72). The initial conformation of the C_2_AB fragment was generated starting from the coordinates of C_2_AB used for the Sqscff system and, after solvation, minimization and equilibration, a 5 ns production MD simulation was performed with position restraints to keep the C_2_B domain at its original position (force constant 1,000 kJ/mol/nm^2^) and additional position restraints to move the C_2_A domain so that its Ca^2+^-binding loops can readily interact with the flat bilayer while the C_2_B domain binds to the flat bilayer through the polybasic face (force constant 100 kJ/mol/nm^2^). The four trans-SNARE complexes generated for the qscv system were used as starting point to generate the four primed complexes. Four Cpx1(27-72) molecules were rotated and translated to interact each with a corresponding trans-SNARE complex based on the crystal structure (Chen et al., 2002), while four C_2_AB molecules were placed at their final designed places. The system was solvated, minimized and equilibrated, and a 1 ns MD simulation was carried out with the following position restraints: i) strong restraints (force constant 4,000 kJ/mol/nm^2^) to keep the synaptobrevin TM regions in their initial positions, as they were intended to remain inserted in the same positions in the vesicle; ii) mild position restraints (force constant 100 kJ/mol/nm^2^) to keep the C_2_AB molecules at their initial (designed) positions; iii) mild position restraints (force constant 100 kJ/mol/nm^2^) on the syntaxin-1 TM regions to move them to their intended designed positions in the flat bilayer, which we planned to place further from the vesicle that in the qscv system; and iv) mild position restraints (force constant 100 kJ/mol/nm^2^) on the N-terminal half of the SNARE four-helix bundle and on Cpx1(27-72) to move them to their designed positions so that each SNARE complex interacted with a corresponding C_2_AB molecule through the primary interface as in the crystal structure (Zhou et al., 2015). Note that we did not include any position restraints on the C-terminal half of the SNARE four-helix bundle and the juxtamembrane regions to allow them to adapt to the imposed restraints. The four resulting primed complexes were merged with the equilibrated vesicle from the qscv system and a square flat bilayer of 31.5 x 31.5 nm^2^, which was placed below the vesicle with a minimum separation of 2.3.

*Second simulation of primed complexes bridging a vesicle and a flat bilayer* (prs2 system). The four primed complexes were generated using the initial primed complexes from the prsg system and running a 50 ns MD simulation with the same restraints used to create the initial complexes. The four resulting primed complexes were merged with the equilibrated vesicle from the qscv system and a flat bilayer of 35 x 35 nm^2^.

*Four C_2_AB molecules bound to Ca^2+^ and to four trans-SNARE complexes bridging a vesicle and a flat bilayer* (prsncpxca system). The Cpx1(27-72) molecules were removed from the four initial primed complexes of the prsg system and five Ca^2+^ ions were added to the corresponding binding sites of each C_2_AB molecule. The resulting complexes were merged with the equilibrated vesicle from the qscv system and the same flat bilayer of 35 x 35 nm^2^ used for the prs2 system.

*Four Ca^2+^-bound C_2_AB molecules not bound to four trans-SNARE complexes bridging a vesicle and a flat bilayer* (cac2absc system). The initial equilibrated prsg system was used as starting point. The Cpx1(27-72) molecules were removed, the C_2_AB molecules were manually moved to areas between the trans-SNARE complexes near but not in contact with the vesicle or the flat bilayer, and five Ca^2+^ ions were added to the corresponding binding sites of each C_2_AB molecule.

### MD simulations

After energy minimization, all systems were heated to 310 K over the course of a 1 ns MD simulation in the NVT ensemble and equilibrated for 1 ns in the NPT ensemble using isotropic Parrinello-Rahman pressure coupling (Stradley et al., 1990). NPT production MD simulations were performed for the times indicated in Table 1 for each system using 2 fs steps, isotropic Parrinello-Rahman pressure coupling and a 1.1 nm cutoff for non-bonding interactions. All simulations were performed at 310 K except one simulation with the qscff system, which was performed at 325 K after a 310 K simulation. Nose-Hoover temperature coupling (Bruch et al., 1992) was used separately for three groups: i) protein atoms plus Ca^2+^ ions if present; ii) lipid atoms; and ii) water and KCL. Periodic boundary conditions were imposed with Particle Mesh Ewald (PME) (Bienstock et al., 1993) summation for long-range electrostatics. The speeds of the production simulations ran on Frontera at TACC are indicated in Table 1.

### Analysis of vesicle-flat bilayer contacts

Because the vesicle and the flat bilayers of the different systems contain large numbers of atoms, it was impractical to analyze vesicle-flat bilayer contacts through measurement of the distances between all atoms of the vesicle and all the atoms of the flat bilayer in multiple time frames of a trajectory. To limit the calculations, we selected only oxygen atoms from frames taken at 1 ns steps of each trajectory and measured the distances between all the oxygen atoms of the vesicle and all the oxygen atoms of the flat bilayer in each frame. The number of vesicle-flat bilayer contacts in each frame was defined as the number of oxygen-oxygen distances below 1 nm.

## Acknowledgments

Most of the work presented in this paper was performed through a Pathways allocation for high performance computing using Frontera (project MCB20033) at the Texas Advanced Computing Center (TACC) at The University of Texas at Austin (URL: http://www.tacc.utexas.edu). This research also used computational resources provided by the BioHPC supercomputing facility located in the Lyda Hill Department of Bioinformatics, UT Southwestern Medical Center, TX (URL: https://portal.biohpc.swmed.edu). This work was supported by grant I-1304 from the Welch Foundation (to JR), by NIH Research Project Award R35 NS097333 (to JR), by NSF Research Project Award MCB-2111728 (to WI), and by the Natural Science Foundation of Shanghai Grant 19ZR1473600 (to YQ).

## Data availability

The files corresponding to our molecular dynamics simulations are being deposited in the dryad database and will be publically available once the deposition is complete.

## Supplementary figures

**Figure 1-figure supplement 1.**
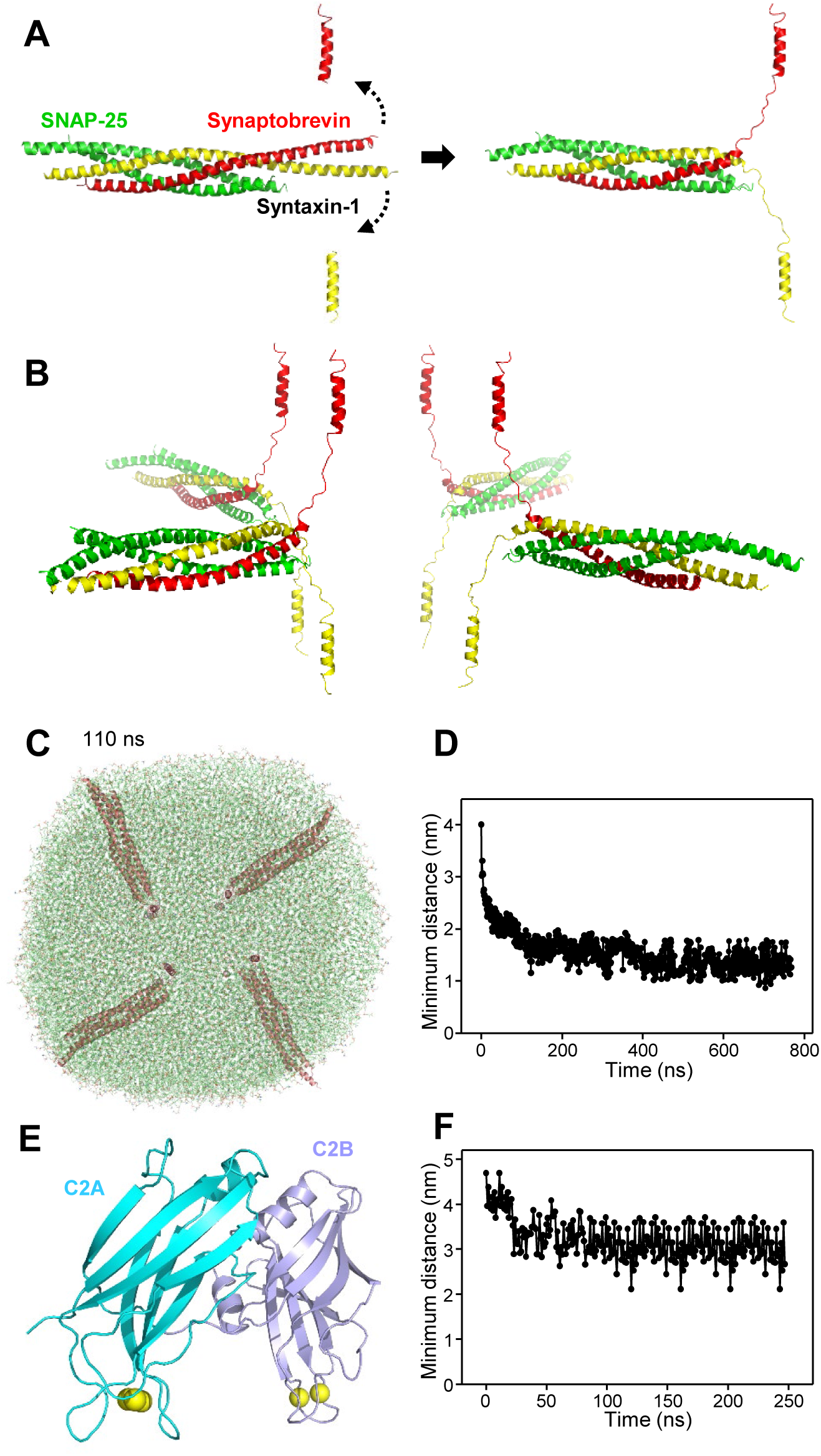
(**A**) Illustration of the procedure used to general the initial structure of a trans-SNARE complex between two flat bilayers. A ribbon diagram of a crystal structure of the neuronal SNARE complex including the TM regions of synaptobrevin and syntaxin-1 (PDB accession code 3HD7) is shown in the middle on the left, with synaptobrevin in red, syntaxin-1 in yellow and SNAP-25 in green. Ribbon diagrams above and below show the positions designed for the TM regions. A 1 ns MD simulation with restraints to force these positions of the TM regions and additional restraints to keep the heavy atoms of the N-terminal half of the SNARE four-helix bundle (up to the polar layer) at their initial positions led to the structure illustrated by the ribbon diagram on the right. (**B**) Ribbon diagrams of the four trans-SNARE complexes generated for the system with two flat bilayers. Three copies of the original structure obtained by the 1 ns restrained MD simulation were generated and then rotated and translated to yield this final configuration. (**C**) Snapshot of the MD simulation of four trans-SNARE complexes between two flat bilayers after 110 ns viewed from the top to illustrate that the flat bilayers acquired an almost circular shape. The SNARE complexes are illustrated by ribbon diagrams in salmon. The lipids are shown as thin stick models. The atom color code for the lipids is: carbon lime, oxygen red, nitrogen blue, phosphorous orange. (**D**) Minimum distance between atoms of the two flat bilayers in frames taken every 1 ns in the simulation of four trans-SNARE complexes between two bilayers. (**E**) Ribbon diagram of the conformation of synaptotagmin-1 C_2_AB molecules used for the simulation of four trans-SNARE complexes and four C_2_AB molecules between two bilayers. The C_2_A domain is colored in cyan and the C_2_B domain in violet. Ca^2+^ ions are shown as yellow spheres. (**F**) Minimum distance between atoms of the two flat bilayers in frames taken every 1 ns in the simulation of four trans-SNARE complexes and four C_2_AB molecules between two bilayers.

**Figure 2-figure supplement 1.**
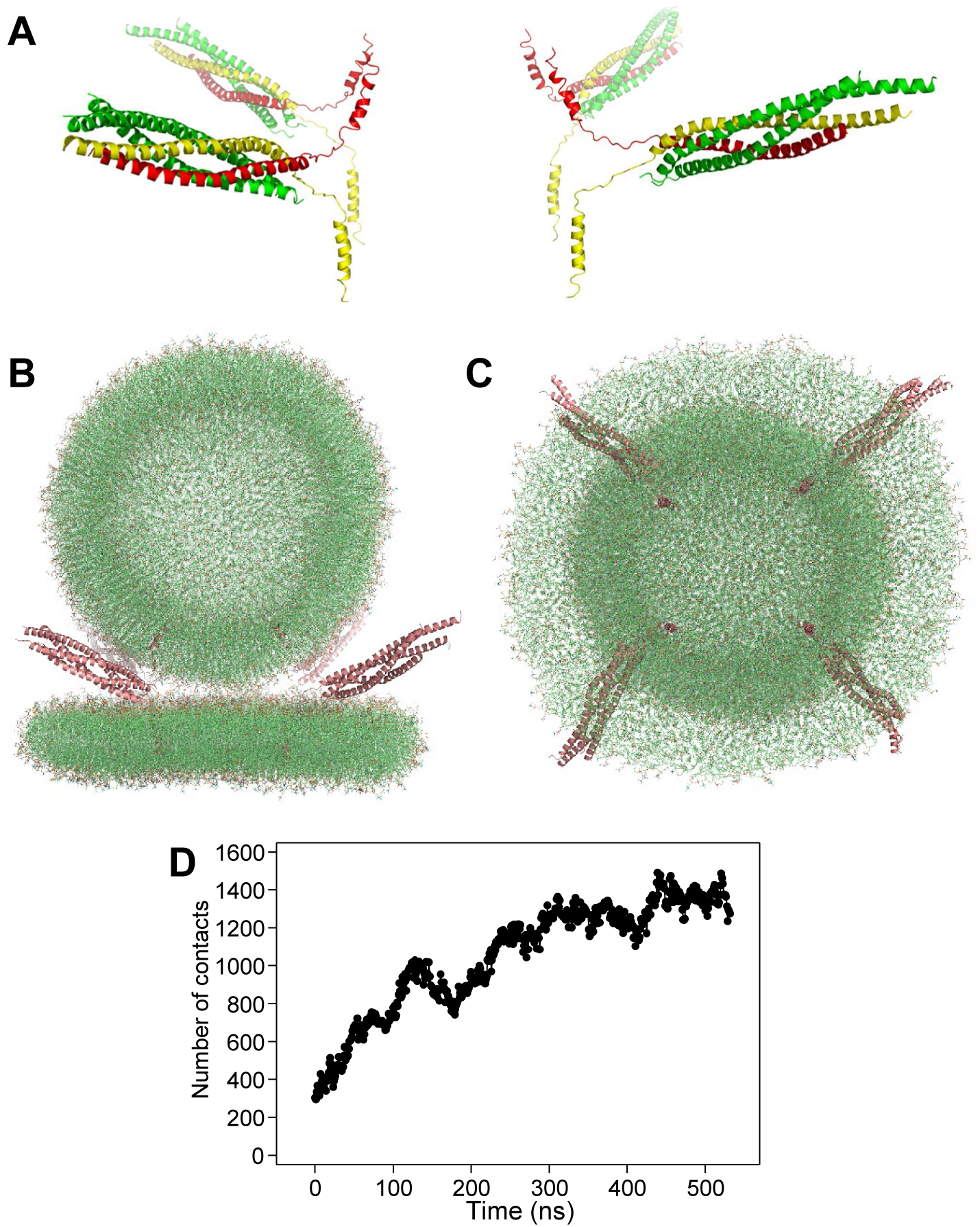
(**A**) Ribbon diagrams of the four trans-SNARE complexes generated for the initial system with one vesicle and a flat bilayer, with synaptobrevin in red, syntaxin-1 in yellow and SNAP-25 in green. (**B,C**) The system of four trans-SNARE complexes bridging a vesicle and a flat bilayer after the equilibration steps viewed from the side (**B**) or from the bottom (**C**). The SNARE complexes are illustrated by ribbon diagrams in salmon. The lipids are shown as thin stick models (carbon lime, oxygen red, nitrogen blue, phosphorous orange). (**D**) Number of contacts in frames taken at 1 ns steps in the simulation of four trans-SNARE complexes bridging a vesicle and a flat bilayer. The number of contacts was defined as the number of distances between oxygen atoms of the vesicle and oxygen atoms of the flat bilayer that were smaller than 1 nm.

**Figure 3-figure supplement 1.**
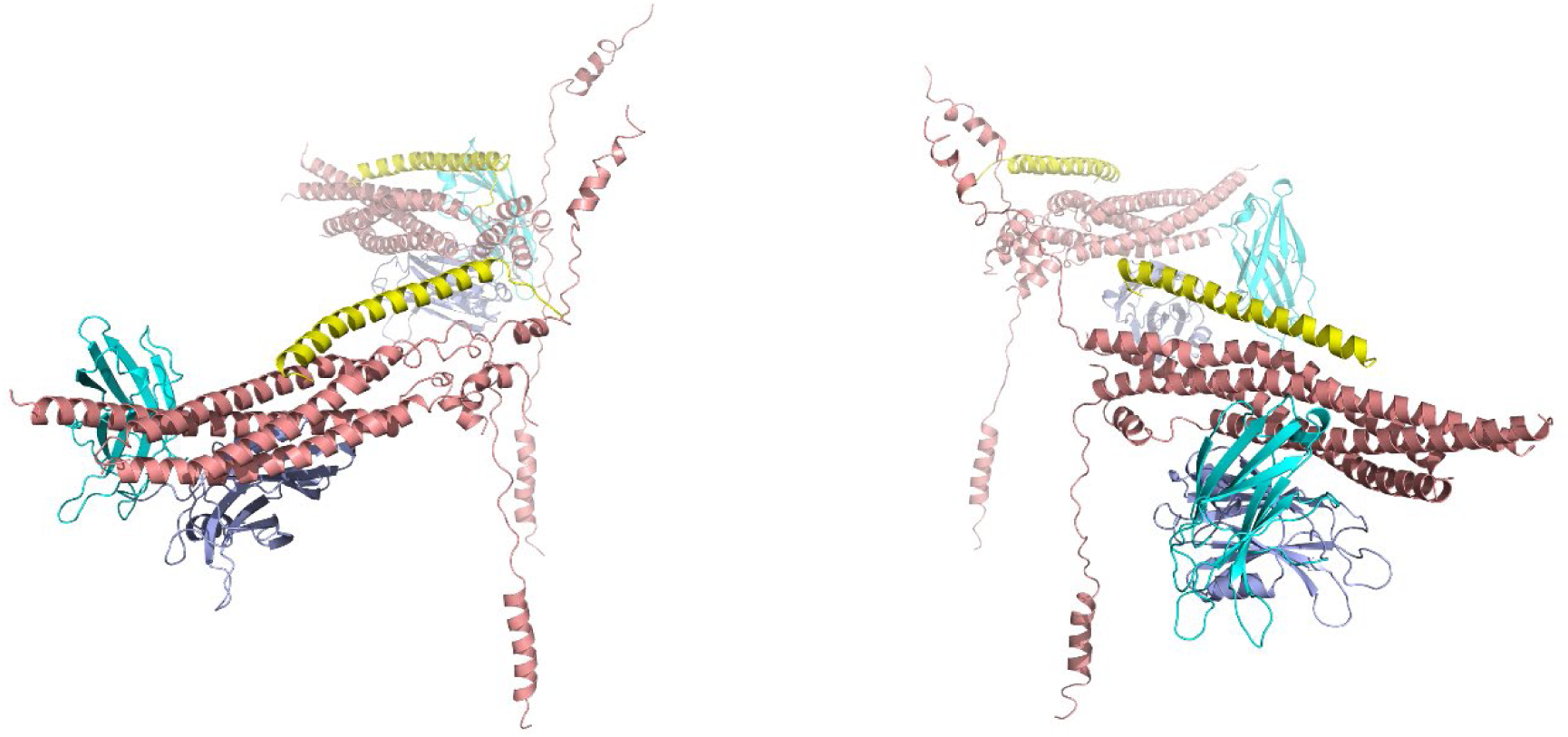
Ribbon diagrams of the four primed complexes generated for the first primed system with one vesicle and a flat bilayer. The SNAREs are in salmon, Cpx1(27-72) in yellow and the synaptotagmin-1 C_2_AB fragment in cyan (C_2_A domain) and violet (C_2_B domain).

**Figure 3-figure supplement 2.**
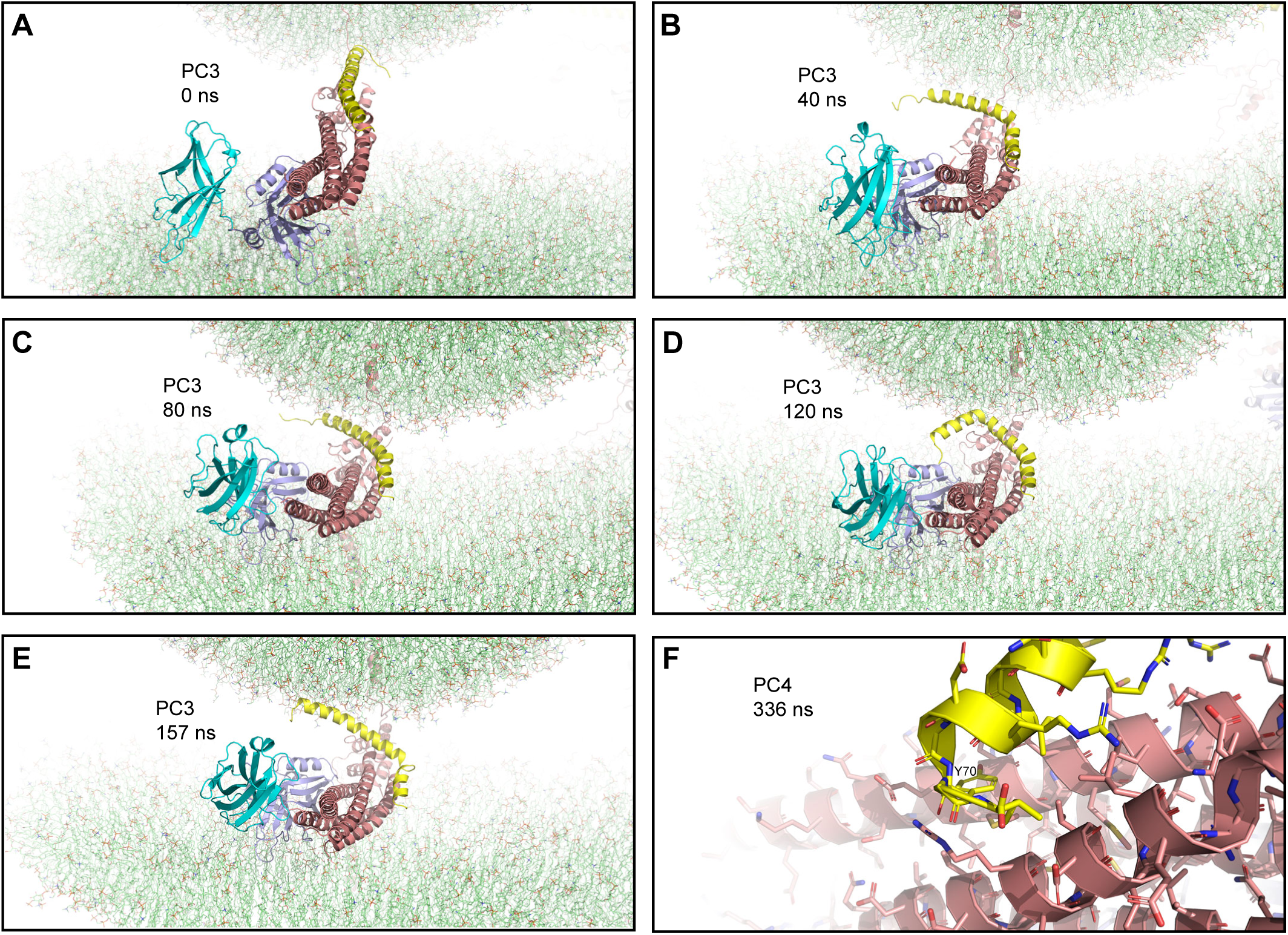
(**A-E**) Close up views of PC3 in the first MD simulation of primed complexes bridging a vesicle and a flat bilayer in the initial configuration (**A**) and after 40, 80, 120 and 157 ns (**B-E**, respectively). PC3 is illustrated by ribbon diagrams, with the SNAREs in salmon, Cpx1(27-72) in yellow and the synaptotagmin-1 C_2_AB fragment in cyan (C_2_A domain) and violet (C_2_B domain). The lipids are shown as thin stick models (carbon lime, oxygen red, nitrogen blue, phosphorous orange). (**F**) Close up view of the region where Cpx1(27-72) remains bound to the SNARE complex in PC4 after 336 ns. Cpx1(27-72) and the SNARE complex are illustrated by ribbon diagrams and stick models with oxygen atoms in red, nitrogen atoms in blue, sulfur atoms in light orange and carbon atoms in yellow [for Cpx1(27-72)] or salmon (for the SNAREs). The position of the Y70 side chain of Cpx1(27-72), which binds at a hydrophobic pocket of the SNARE complex, is indicated.

**Figure 4-figure supplement 1.**
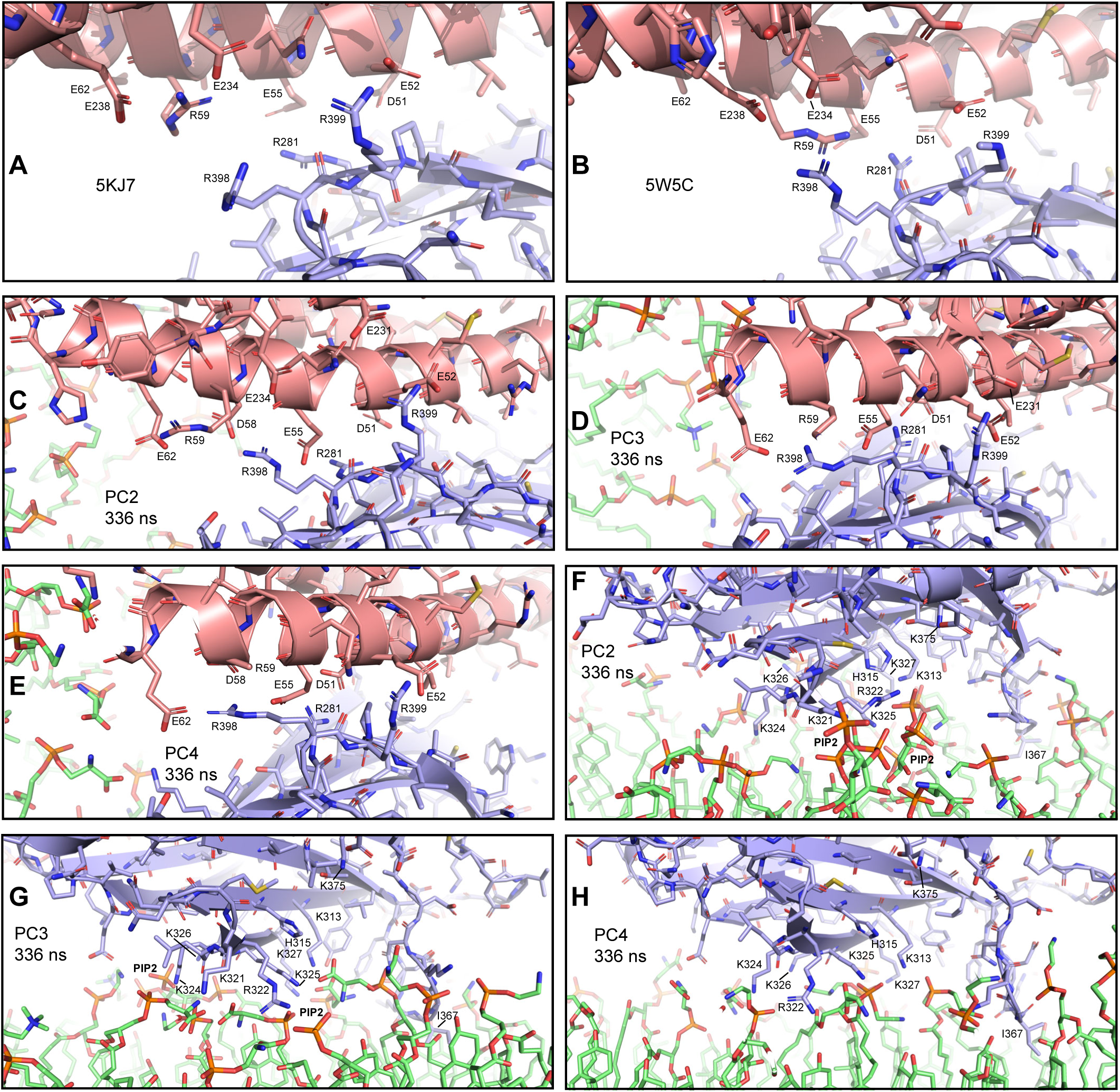
(**A-E**) Close up views of site II of the primary interface between the C_2_B domain and the SNARE complex in two crystal structures (**A**, PDB accession number 5KJ7; **B**, PDB accession number 5W5C), and in PC2, PC3 and PC4 (**C-E**, respectively) after 336 ns of the first MD simulation of primed complexes bridging a vesicle and a flat bilayer. The C_2_B domain and the SNARE complex are illustrated by ribbon diagrams and stick models with oxygen atoms in red, nitrogen atoms in blue, sulfur atoms in light orange and carbon atoms in violet [for the C_2_B domain] or salmon (for the SNAREs). The positions of selected side chains are indicated. (**F-H**) Close up views of the interaction of the C_2_B domain of PC2, PC3 or PC4 with the flat bilayer after 336 ns (**F-H**, respectively). The positions of PIP_2_ headgroups, basic side chains involved in interactions with the lipids, and the hydrophobic side chain of I367 at the tip of a Ca^2+^-binding loop that inserts into the bilayer, are indicated.

**Figure 4-figure supplement 2.**
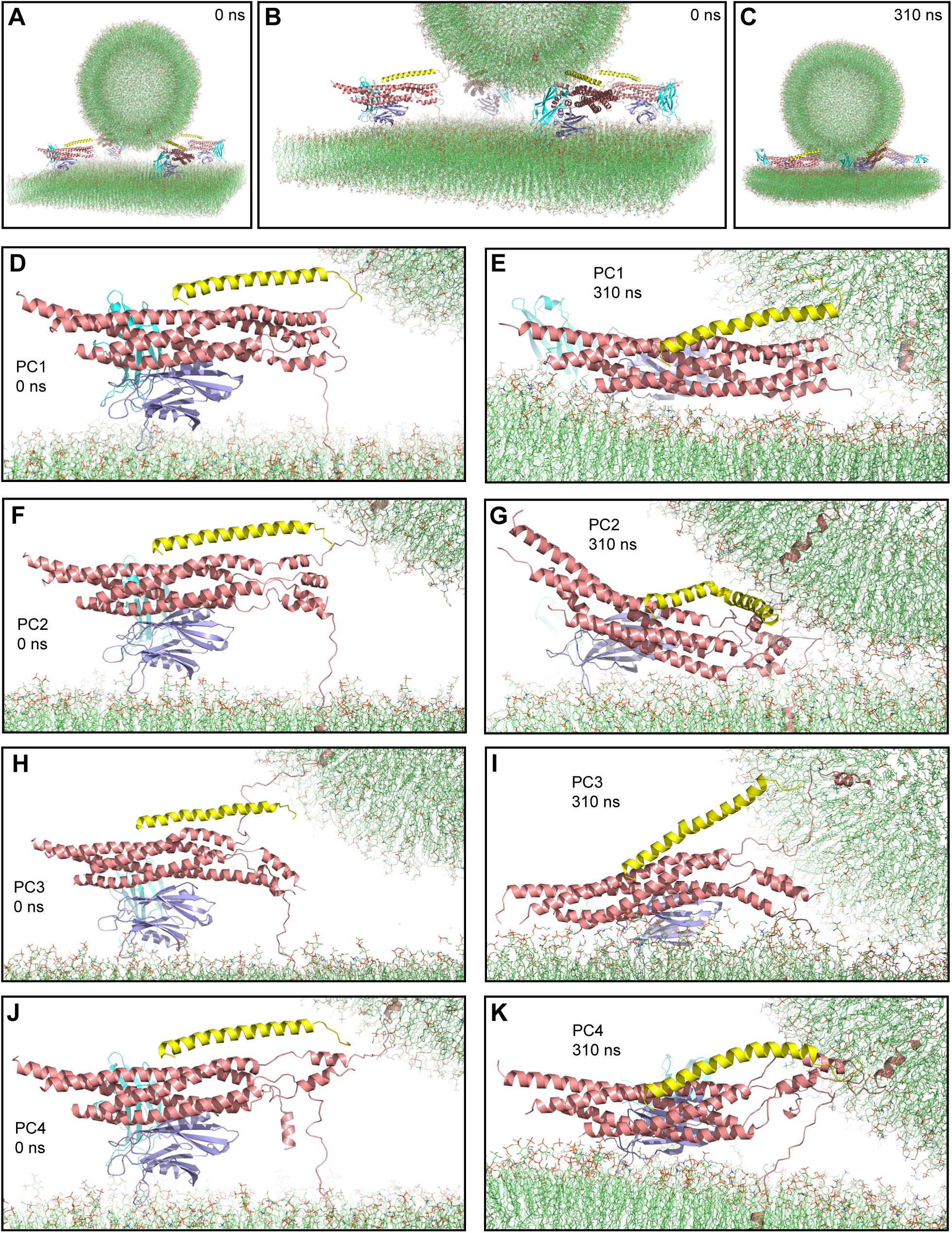
Second MD simulation of primed complexes bridging a vesicle and a flat bilayer. (**A**) Overall view of the initial system. (**B**) Close up view of the four primed complexes in the initial system. (**C**) Snapshot of the system after a 310 ns MD simulation. (**D-K**) Close up views of the individual primed complexes (named PC1-PC4) in the initial configuration (**D,F,H,J**) and after the 310 ns MD simulation (**E,G,I,K**). In all panels, the primed complexes are illustrated by ribbon diagrams, with the SNAREs in salmon, Cpx1(27-72) in yellow and the synaptotagmin-1 C_2_AB fragment in cyan (C_2_A domain) and violet (C_2_B domain). The lipids are shown as thin stick models (carbon lime, oxygen red, nitrogen blue, phosphorous orange).

**Figure 4-figure supplement 3.**
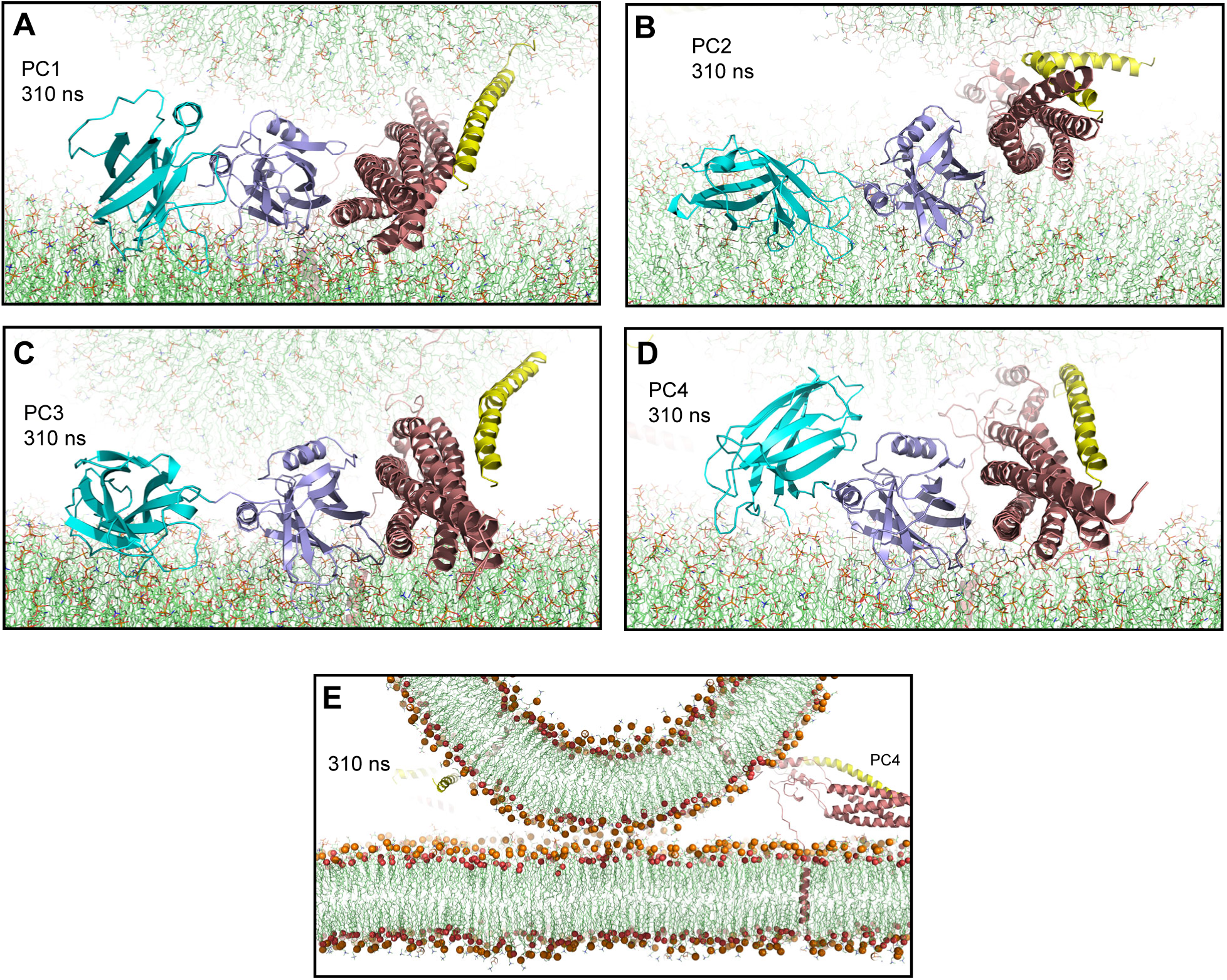
Additional views of the second MD simulation of primed complexes bridging a vesicle and a flat bilayer. (**A-D**) Close up views of the four primed complexes after 310 ns showing how the synaptotagmin-1 C_2_B domain binds to the SNARE complex through the primary interface and to the flat bilayer with the polybasic face, which dictates that the Cpx1(27-72) helix is oriented toward the vesicle and bends in different ways and directions to avoid steric clashes. This arrangement forces the SNARE four-helix bundle to be close to the flat bilayer. The primed complexes are illustrated by ribbon diagrams, with the SNAREs in salmon, Cpx1(27-72) in yellow and the synaptotagmin-1 C_2_AB fragment in cyan (C_2_A domain) and violet (C_2_B domain). The lipids are shown as thin stick models (carbon lime, oxygen red, nitrogen blue, phosphorous orange). (**E**) Thin slice of the system showing a point-of-contact interface between the vesicle and the flat bilayer at 310 ns. Phosphorous atoms of phospholipids and the oxygen atoms of cholesterol molecules are shown as spheres to illustrate the approximate locations of lipid head groups. The positions of PC1 and PC3 are indicated.

**Figure 4-figure supplement 4.**
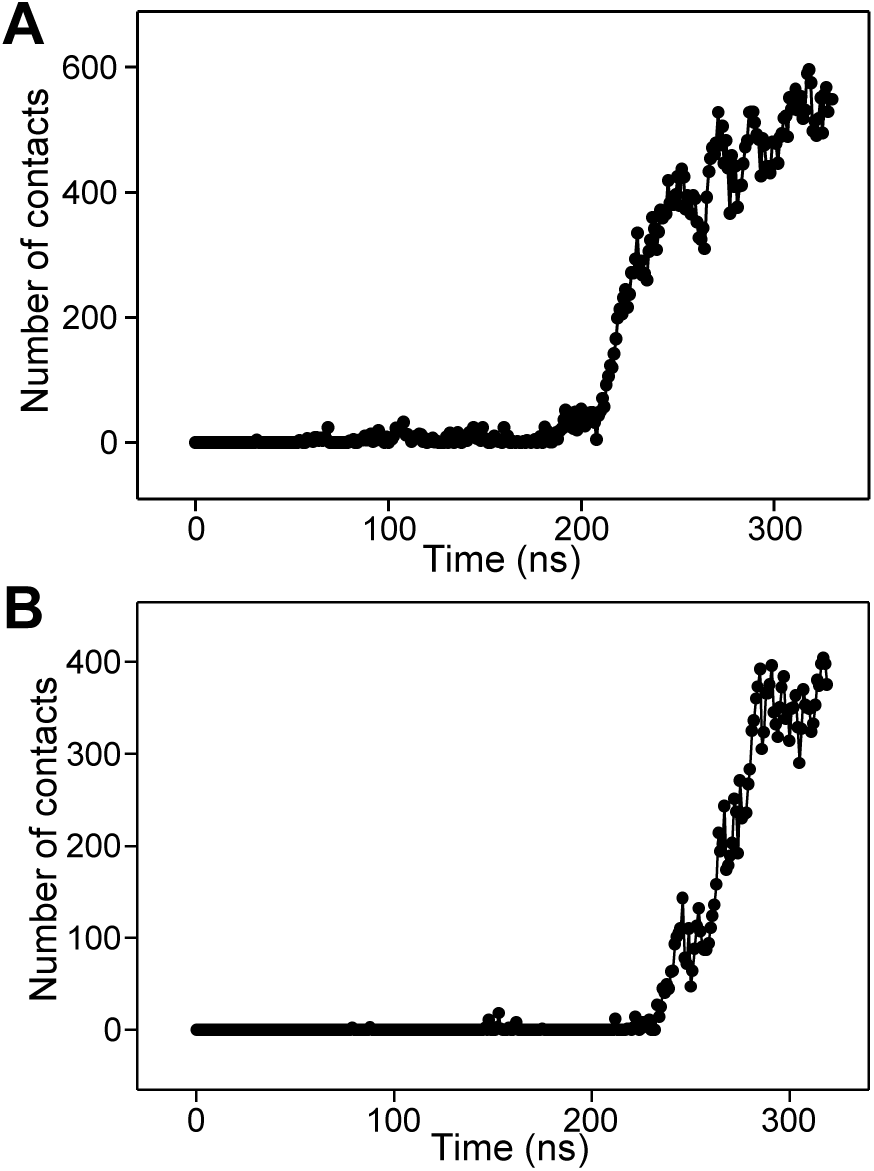
Number of contacts in frames taken at 1 ns steps in the first (**A**) or second (**B**) simulation of four primed complexes bridging a vesicle and a flat bilayer. The number of contacts was defined as the number of distances between oxygen atoms of the vesicle and oxygen atoms of the flat bilayer that were smaller than 1 nm.

**Figure 5-figure supplement 1.**
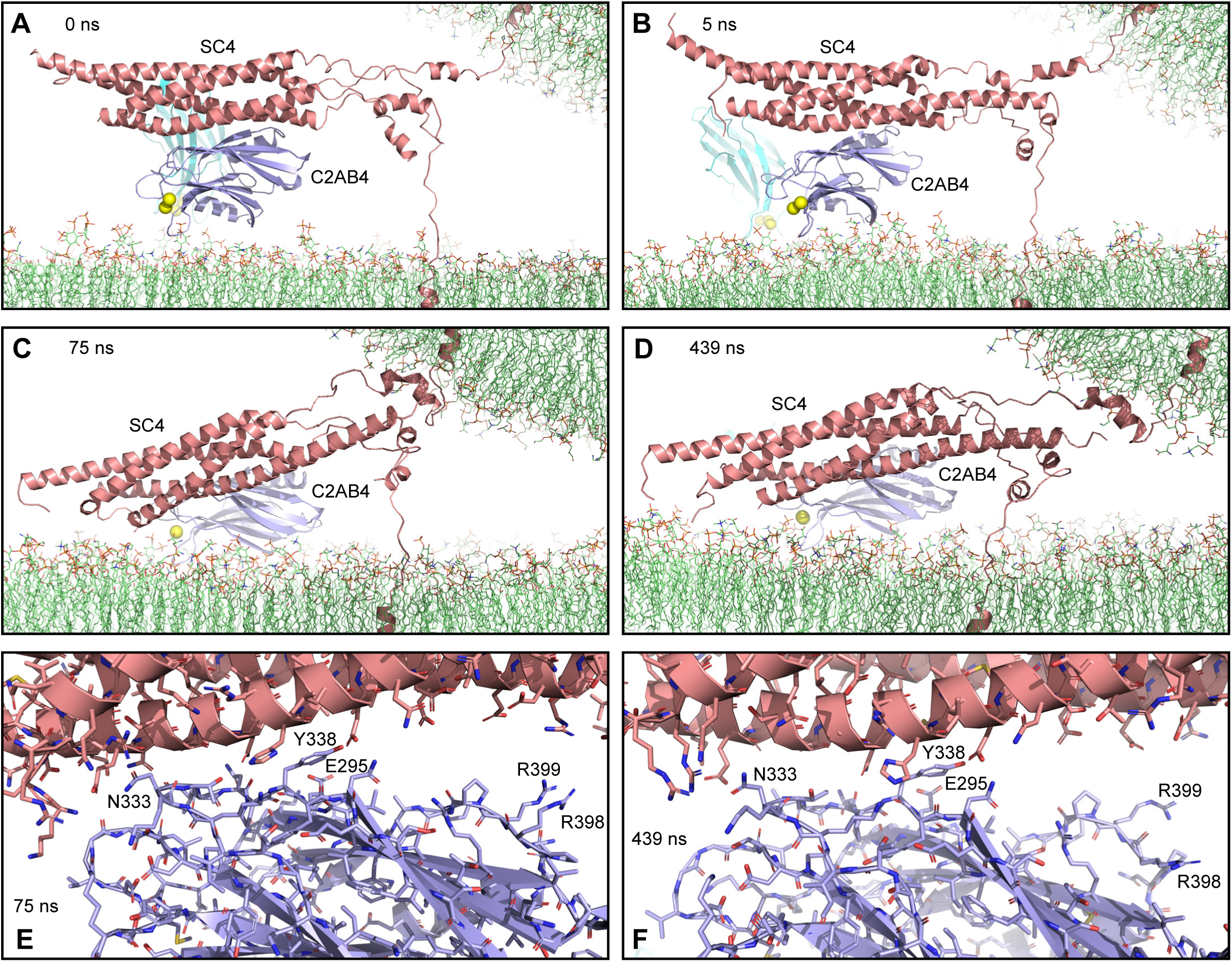
(**A-D**) Close up views of SC4 and C2AB4 in the MD simulation of C_2_AB bound to Ca^2+^ and to trans-SNARE complexes bridging a vesicle and a flat bilayer in the initial configuration (**A**) and after 5, 75 and 439 ns (**B-D**, respectively). The SNAREs are represented by ribbon diagrams in salmon and the synaptotagmin-1 C_2_AB fragment by ribbon diagrams in cyan (C_2_A domain) and violet (C_2_B domain). Ca^2+^ ions are shown as yellow spheres. The lipids are shown as thin stick models (carbon lime, oxygen red, nitrogen blue, phosphorous orange). Note that in just the first 5 ns one of the helices that was disrupted was almost fully formed but overall there was no substantial progress toward assembly of the C-terminal part of the SNARE four-helix bundle in 439 ns. (**E-F**) Close up views of the primary interface between SC4 and C2AB4 after 75 (**E**) and 439 (**F**) ns. The C_2_B domain and the SNARE complex are illustrated by ribbon diagrams and stick models with oxygen atoms in red, nitrogen atoms in blue, sulfur atoms in light orange and carbon atoms in violet [for the C_2_B domain] or salmon (for the SNAREs). The positions of selected side chains are indicated.

**Figure 5-figure supplement 2.**
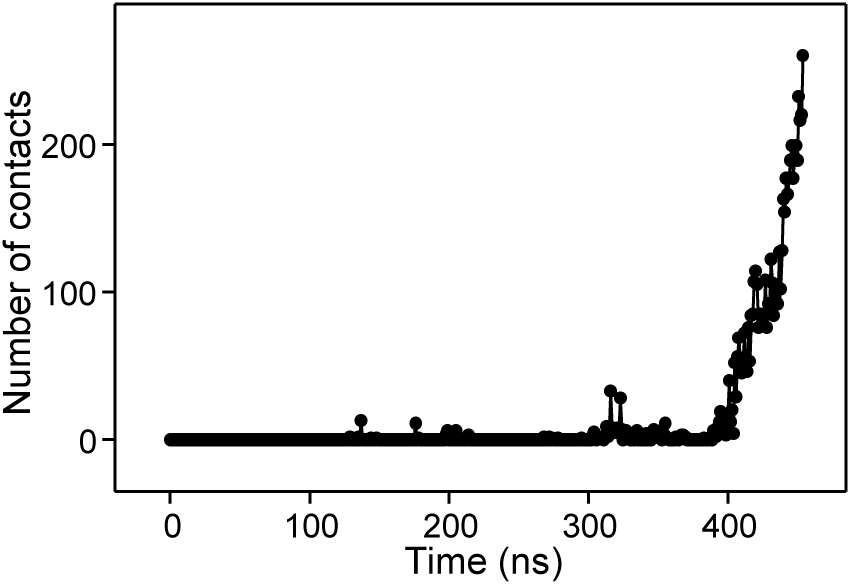
Number of contacts in frames taken at 1 ns steps in the MD simulation of C_2_AB bound to Ca^2+^ and to trans-SNARE complexes bridging a vesicle and a flat bilayer. The number of contacts was defined as the number of distances between oxygen atoms of the vesicle and oxygen atoms of the flat bilayer that were smaller than 1 nm.

**Figure 6-figure supplement 1.**
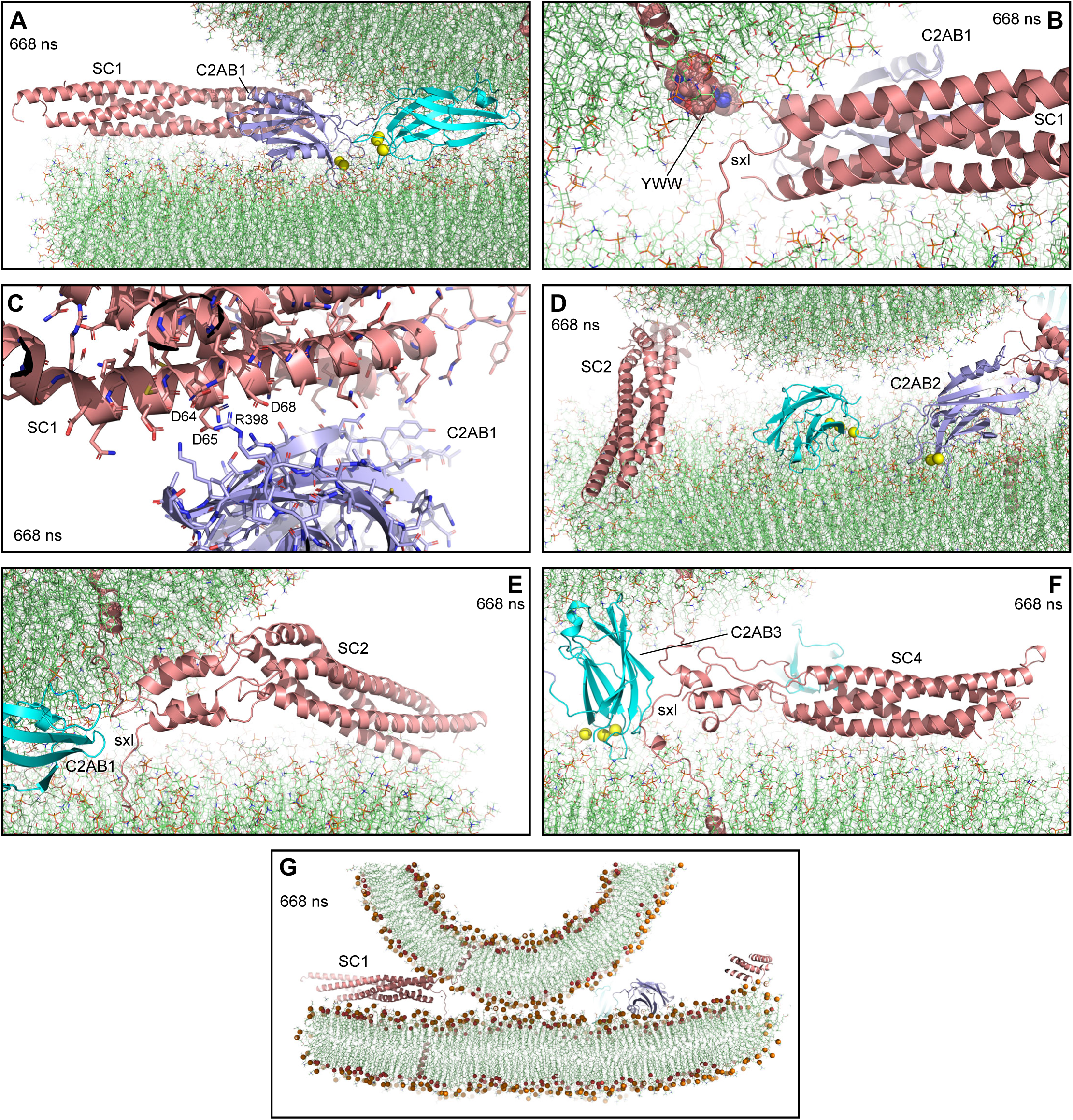
MD simulation of Ca^2+^-bound C_2_AB dissociated from trans-SNARE complexes bridging a vesicle and a flat bilayer. (**A-B**) Close up views from different angles of C2AB1 and SC1 after 668 ns. The SNAREs are represented by ribbon diagrams in salmon and the synaptotagmin-1 C_2_AB fragment by ribbon diagrams in cyan (C_2_A domain) and violet (C_2_B domain). Ca^2+^ ions are shown as yellow spheres. The lipids are shown as thin stick models (carbon lime, oxygen red, nitrogen blue, phosphorous orange). Panel (**A**) shows how the Ca^2+^-binding loops of the two C_2_ domains point toward the center of the vesicle-flat bilayer interface, next to the TM regions of synaptobrevin and syntaxin-1. Panel (**B**) shows how the three aromatic residues from the synaptobrevin juxtamembrane linker (Tyr88, Trp89 and Trp90, abbreviated YWW and shown as spheres) are inserted into the vesicle lipid bilayer, and part of the syntaxin-1 juxtamembrane linker (sxl) is not inserted into the flat bilayer. (**C**) Close up view of the interface between the C_2_B domain of C2AB1 and SC1 after 668 ns. The C_2_B domain and the SNARE complex are illustrated by ribbon diagrams and stick models with oxygen atoms in red, nitrogen atoms in blue, sulfur atoms in light orange and carbon atoms in violet [for the C_2_B domain] or salmon (for the SNAREs). The positions of selected side chains are indicated. (**D-F**) Close up views of SC2 from different angles (**D-E**) and of SC4 (**F**), showing the relative positions of SC2 and SC4 with respect to nearby C_2_AB molecules. Molecules are represented as in (**A**). (**G**) Thin slice of the system showing a point-of-contact interface between the vesicle and the flat bilayer at 668 ns. Phosphorous atoms of phospholipids and the oxygen atoms of cholesterol molecules are shown as spheres to illustrate the approximate locations of lipid head groups. The position of SC1 is indicated.

**Figure 6-figure supplement 2.**
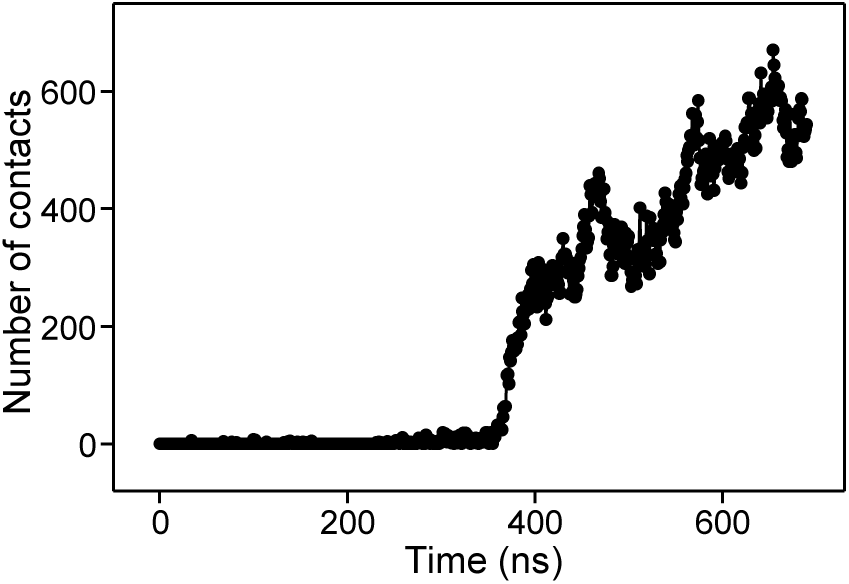
Number of contacts in frames taken at 1 ns steps in the simulation of Ca^2+^-bound C_2_AB dissociated from trans-SNARE complexes bridging a vesicle and a flat bilayer. The number of contacts was defined as the number of distances between oxygen atoms of the vesicle and oxygen atoms of the flat bilayer that were smaller than 1 nm.

## Notes

### Competing Interest Statement

The authors have declared no competing interest.

